# Comparison of Wild-Type and High-risk PNPLA3 variants in a Human Biomimetic Liver Microphysiology System for Metabolic Dysfunction-associated Steatotic Liver Disease Precision Therapy

**DOI:** 10.1101/2024.04.22.590608

**Authors:** Mengying Xia, Mahboubeh Varmazyad, Iris Pla-Palacín, Dillon C. Gavlock, Richard DeBiasio, Gregory LaRocca, Celeste Reese, Rodrigo Florentino, Lanuza A.P. Faccioli, Jacquelyn A. Brown, Lawrence A. Vernetti, Mark Schurdak, Andrew M. Stern, Albert Gough, Jaideep Behari, Alejandro Soto-Gutierrez, D. Lansing Taylor, Mark T. Miedel

## Abstract

Metabolic dysfunction-associated steatotic liver disease (MASLD) is a worldwide health epidemic with a global occurrence of approximately 30%. The pathogenesis of MASLD is a complex, multisystem disorder driven by multiple factors including genetics, lifestyle, and the environment. Patient heterogeneity presents challenges for developing MASLD therapeutics, creation of patient cohorts for clinical trials and optimization of therapeutic strategies for specific patient cohorts. Implementing pre-clinical experimental models for drug development creates a significant challenge as simple *in vitro* systems and animal models do not fully recapitulate critical steps in the pathogenesis and the complexity of MASLD progression. To address this, we implemented a precision medicine strategy that couples the use of our liver acinus microphysiology system (LAMPS) constructed with patient-derived primary cells. We investigated the MASLD-associated genetic variant PNPLA3 rs738409 (I148M variant) in primary hepatocytes, as it is associated with MASLD progression. We constructed LAMPS with genotyped wild type and variant PNPLA3 hepatocytes together with key non-parenchymal cells and quantified the reproducibility of the model. We altered media components to mimic blood chemistries, including insulin, glucose, free fatty acids, and immune activating molecules to reflect normal fasting (NF), early metabolic syndrome (EMS) and late metabolic syndrome (LMS) conditions. Finally, we investigated the response to treatment with resmetirom, an approved drug for metabolic syndrome-associated steatohepatitis (MASH), the progressive form of MASLD. This study using primary cells serves as a benchmark for studies using “patient biomimetic twins” constructed with patient iPSC-derived liver cells using a panel of reproducible metrics. We observed increased steatosis, immune activation, stellate cell activation and secretion of pro-fibrotic markers in the PNPLA3 GG variant compared to wild type CC LAMPS, consistent with the clinical characterization of this variant. We also observed greater resmetirom efficacy in PNPLA3 wild type CC LAMPS compared to the GG variant in multiple MASLD metrics including steatosis, stellate cell activation and the secretion of pro-fibrotic markers. In conclusion, our study demonstrates the capability of the LAMPS platform for the development of MASLD precision therapeutics, enrichment of patient cohorts for clinical trials, and optimization of therapeutic strategies for patient subgroups with different clinical traits and disease stages.

## Introduction

Metabolic-dysfunction associated steatotic liver disease (MASLD), previously named non-alcoholic fatty liver disease (NAFLD), is a chronic liver disease affecting ∼ 30% of the global adult population (1–4). Management of MASLD presents a huge financial burden on health care systems today affecting 30 million adults in the US which is expected to increase to over 100 million by 2030 (5). MASLD is a complex, heterogeneous disease involving liver metabolic dysfunction associated with possible co-morbidities including T2D, obesity and hypertension, with further heterogeneity from genetic risk factors (6–8), as well as patient lifestyle and environment. The progressive form of MASLD, metabolic-dysfunction associated steatohepatitis (MASH), is a major cause of hepatocellular carcinoma and is also the leading indication for liver transplantation in the United States (9).

The last decade has produced a major effort in developing MASLD/MASH therapeutics with approximately 1,300 currently registered MASLD/MASH clinical trials (https://clinicaltrials.gov) (10, 11). The high failure rate in developing therapeutics for MASLD patients is due to a combination of factors including a) the complex, heterogeneous nature of the disease, including a variety of possible co-morbidities; b) genotypic differences in key MASLD-associated genes; c) patient lifestyle and environment; as well as d) the historical reliance on animal models of disease that do not fully recapitulate the human disease complexity and heterogeneity (12–17). Recently, resmetirom (Rezdiffra™), became the first approved drug for MASH with stage 2 or 3 fibrosis; however, it only impacts approximately 25% of the patients treated in clinical trials (18). The complex and heterogeneous nature of MASLD progression suggests that a precision medicine approach is required for success in developing therapeutics, segmenting MASLD patients into specific cohorts for enrollment in clinical trials and optimizing therapeutic strategies for specific patient cohorts. A critical question for clinicians is how to predict responders more accurately from non-responders for any given MASLD therapeutic. We implemented a precision medicine approach harnessing the use of genotyped, human primary cells and patient-derived induced pluripotent stem cells (iPSCs) differentiated into the major liver cell types to construct patient-specific MPS. We investigated the combined role of a key MASLD-associated genetic variant, as well as eating lifestyle by altering media formulations to model different blood chemistries to define their roles in disease severity and progression and response to resmetirom (**Figure 1)**.

**Figure 1.**
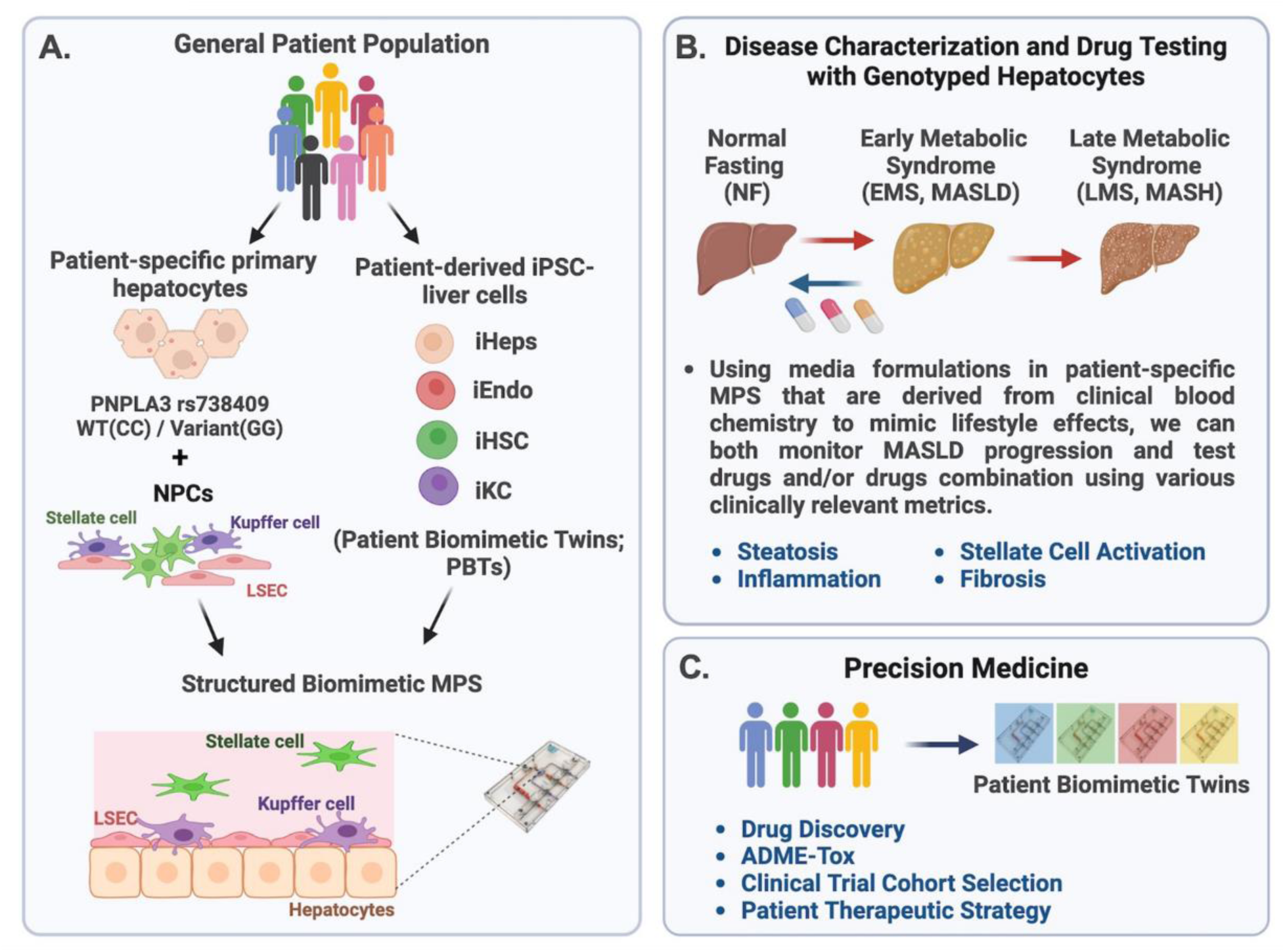
Overview of the precision medicine approach to investigate genotype-specific MASLD progression and drug response using PNPLA3 genotyped patient-derived cells in the liver acinus microphysiology system (LAMPS). (A) MASLD progression is complex due to multiple factors including genetics, environment, and lifestyle, resulting in patient heterogeneity. The PNPLA3 rs738409 GG variant is highly associated with MASLD susceptibility (23, 24, 98, 99). To study this high-risk variant, we started with genotyped PNPLA3 rs738409 GG variant and CC wild type primary hepatocytes to use as a benchmark for studies using iPSC-derived liver cells (100, 101). These genotyped primary hepatocytes were cultured with nonparenchymal cells (NPCs) including liver sinusoidal endothelial cells (LSECs), stellate cells, and Kupffer cells in LAMPS (46, 47, 63). (B) MASLD progression in LAMPS was driven with specific media formulations to mimic lifestyle effects as previously described (47), and monitored through various clinically relevant metrics, including steatosis, stellate cell activation, inflammation, and fibrosis. Additionally, the efficacy of recently FDA-approved resmetirom (Rezdiffra^TM^) (18, 87) was assessed in LAMPS, focusing on the early stage of MASLD progression using EMS medium. (C) Results from these studies can then inform a precision medicine strategy that implements the use of patient-derived iPSCs to generate patient biomimetic twins (PBTs) for drug testing and development as well as clinical trial patient cohort selection (40).

Genetic factors contribute to MASLD development and progression (19, 20). A correlation between increased risk factors and MASLD patients carrying specific gene variants, including PNPLA3 rs738409, MBOAT7 rs641738, TM6SF2 rs58542926 and GCKR rs780094 is well known (21, 22). Among these noted genetic variants, evidence shows that the human patatin-like phospholipase domain-containing-3 (PNPLA3) gene rs738409 C>G polymorphism *(PNPLA3 rs738409 / I148M)* has a strong correlation to increased MASLD severity including hepatic steatosis, fibrosis, cirrhosis, and hepatocellular carcinoma (23–29). In vitro experiments show that PNPLA3 has triacylglycerol hydrolase, acyltransferase, and transacylase activities which regulate lipid droplet remodeling in both hepatocytes and hepatic stellate cells (30–32). The PNPLA3 C>G (GG) variant results in a reduction in fatty acid hydrolysis and impaired mobilization of triglycerides, resulting in hepatic triglyceride accumulation. (23, 33). While largely unknown, it has been observed that patients carrying the GG variant show differential response to some drug treatments compared to patients carrying wild type (CC) PNPLA3 (34–36). Thus, this differential response provides an initial basis for stratifying MASLD patients according to their PNPLA3 genotype, offering an initial strategy for implementing a precision medicine-based approach for investigating disease progression and assessing drug responses within specific PNPLA3 cohorts.

Animal models to study MASLD progression and evaluate key disease-associated genetic variants such as the PNPLA3 GG polymorphism have been characterized with various metrics demonstrating increased steatosis, inflammation, fibrogenesis, oxidative stress and insulin resistance, yet do not fully recapitulate the complex and heterogeneous progression of the human disease (12–17). Human microphysiology systems (MPS) are experimental models designed to recapitulate the structure and both normal and disease state physiology of tissues and organs to serve as a complement to existing animal models (37, 38). MPS are 3D microfluidic platforms composed of multiple cell types that mimic overall organ structure and provide cell-to-cell communication using human primary cells, immortalized cell lines, and iPSCs (39–41). Recently, multiple human liver MPS have evolved and been implemented to study mechanisms of MASLD pathogenesis and serve as drug testing platforms (40, 42–60). We have implemented the liver acinus microphysiology system (LAMPS), a structured biomimetic, that is a 3D layered model constructed through a combination of sequential cell layering and cell-to-cell self-organization between 4 key liver cell types including primary hepatocytes and liver sinusoidal endothelial cells (LSECs) as well as cell lines for hepatic stellate cells (the LX-2 cell line) and Kupffer-like cells (THP-1 cell line), and maintained under flow to mimic either Zone 1 or Zone 3 oxygen tensions (61–63) **(Fig S1).** The LAMPS has been tested and reproduced by the Texas A&M Tissue Chip Validation Center (Tex-Val), one of the National Center for Advancing Translational Sciences (NCATS) funded Tissue Chip Testing Centers (TCTC) and has demonstrated reproducible features for hepatic function and (46, 47, 63–69).

Our overall strategy towards implementing a precision medicine platform for MASLD has been to first test the MASLD LAMPS model with key primary cells as a benchmark for the next phase using “patient biomimetic twins” (PBTs) produced using patient, induced pluripotent stem cells (iPSCs) (40) **(Fig 1A).** Using media formulations derived from clinical blood chemistry indicators in genotyped patient-specific primary cell LAMPS or PBTs, we then monitor both MASLD progression and test drug efficacy using a panel of disease-relevant metrics **(Fig 1B)** (46, 47). The results from these studies coupled with the use of patient information and clinomics inform a precision medicine strategy using PBTs for drug testing and development to inform clinical trial cohort selection **(Fig 1C)** (**68**).

In this study, we aimed to determine if the MASLD LAMPS constructed with genotyped PNPLA3 GG variant or CC wild type primary hepatocytes exhibited differences in MASLD progression and drug efficacy at an early stage of the disease. MASLD progression was driven using media formulations to mimic disease progression from normal fasting (NF) to early metabolic syndrome (EMS) and then late metabolic syndrome (LMS) for 8 days (46, 47, 68). Disease progression and response to resmetirom treatment were monitored through multiple clinically relevant metrics including steatosis, inflammation stellate cell activation and fibrosis (18).

## Materials and Methods

### Cell sources and initial culture

Cryopreserved primary human hepatocytes were genotyped for PNPLA3 rs738409(G) using TaqMan® SNP Genotyping Assays (Life Technologies, Assay ID C_7241_10, 4351379) according to the manufacturer’s protocol. PNPLA3 GG homozygotes and CC homozygotes with >90% viability, post-thaw re-plating efficiency > 90%, and suitable for long-term culture were selected and purchased from Discovery Life Sciences (Gentest^®^ 999Elite^™^ Human Hepatocytes; catalog #82006); GG variant lots: HH1142, HH1072; CC wild type lots: HH1136, HH1178 and Thermo Fisher Scientific (Human Plateable Hepatocytes, Hu8391 (CC wild type), catalog # HMCPTS). Human liver sinusoidal endothelial cells (LSECs) were purchased from Life Net Health (NPC-AD-LEC-P1). The human monoblast cell line, THP-1, used to generate Kupffer-like cells, was purchased from ATCC (TIB-202) and LX-2 human stellate cells were purchased from Sigma Aldrich (SCC064). LSECs were cultured in endothelial cell basal medium-2 (EBM-2, Lonza, CC-3162). THP-1 cells were cultured in suspension in RPMI 1640 Media (Cytiva, SH30096.FS) supplemented with 10% Regular Fetal Bovine Serum (FBS; Corning, MT35010CV), 100 μg/mL penicillin streptomycin (Cytiva, SV30010), and 2 mM L-Glutamine (Cytiva, SH30034.01). THP-1 cells were differentiated into mature macrophages by treatment with 200 ηg/mL phorbol myristate acetate (Sigma Aldrich, 524400) for 48 h. LX-2 cells were cultured in Dulbecco’s Modified Eagle Medium (DMEM; Thermo Fisher Scientific, 11965118) supplemented with 2% FBS and 100 units/ml penicillin and 100 μg/mL streptomycin.

### LAMPS assembly and maintenance

LAMPS studies were carried out as previously described for model assembly (40, 46, 47, 61–63) **(Figure S1)**. Briefly, LAMPS models were constructed using the HAR-V single channel device (SCC-001) from Nortis, which composed of four key liver cell types and used at the following cell densities: primary cryopreserved human hepatocytes (2.75 x 10^6^ cells/mL), primary liver sinusoidal endothelial cells (LSECs 1.5 x 10^6^ cells/mL), and THP-1 (0.4 x 10^6^ cells/mL) and LX-2 (0.2 x 10^6^ cells/mL). The percentages of hepatocytes, THP-1, LSEC, and LX-2 cells are consistent with the scaling used in our previously published models [1-5]. The interior of the devices was coated with 100 mg/mL bovine fibronectin (Sigma Aldrich, F1141) and 150 mg/mL rat tail collagen (Corning, 354249) in PBS prior to cell seeding. For all steps involving injection of media and/or cell suspensions into LAMPS devices, 100–150 μl per device was used to ensure complete filling of fluidic pathways, chamber, and bubble traps. The devices were then overlayed with 2.5 mg/ml rat tail collagen I (Corning) and maintained with the perfusion of different conditions for 8 days at a flow rate of 5 μL/h to recapitulate zone 3 oxygen tension (61).

### Formulation of normal fasting and early metabolic syndrome, and late metabolic syndrome media

Media were formulated as previously described [2] to create disease progression from Normal Fasting (NF) to Early Metabolic Syndrome (EMS or MASLD) and Late Metabolic Syndrome (LMS or late stage MASLD). The media were formulated with glucose-free Williams E medium (Thermo Fisher Scientific, ME18082L1) as the base medium supplemented with physiologically relevant levels of glucose (Sigma Aldrich, G8644), insulin (Thermo Fisher Scientific, 12585014), glucagon (Sigma Aldrich, G2044), oleic acid (Cayman Chemicals, 29557), palmitate acid (Cayman Chemicals, 29558) and molecular drivers of disease including TGF-β1 (Thermo Fisher Scientific, PHG9214) and Lipopolysaccharide (Sigma Aldrich, L2654) as shown in **Table S1**.

### 96-well plate LAMPS assembly and maintenance

The 96-well plate LAMPS was used for the lipid peroxidation analysis. The construction of 96-well plate LAMPS was conducted as previously described (62). Briefly, the 96-well plate LAMPS followed the same percentage of each cell type in LAMPS model. Hepatocytes (50,000 cells/well) were plated in a collagen I-coated clear-bottom 96 well plate (Thermo Fisher Scientific, 08-774-307). Porcine liver extracellular matrix (LECM) (400 μg/mL) was added on the top of the hepatocytes to create a thin matrix layer. A mixture of LSEC (0.54 x 10^6^ cells/mL) and THP-1 (0.28 x10^6^ cells/mL) was added on the top of LECM finally an overlay of LX-2 (0.1 x 10^6^ cells/mL) suspended in a 2.5 mg/mL solution of rat-tail collagen (pH 7.2) was added. 96-well plate LAMPS was maintained in 100 μL LAMPS perfusion media. Media were replaced every 48 h during the experimental time course.

### Image and analysis for lipid peroxidation (LPO) level

BODIPY 581/591 C11 (Lipid Peroxidation Sensor, Thermo Fisher Scientific, D3861) was added to the 96-well co-culture model at day 5 with a final concentration of 5 µM in the appropriate medium (NF, EMS, or LMS) and incubated for 30 min at 37°C. Hydrogen peroxide was used as a positive control to induce LPO. Prior to imaging, the cells were washed twice with PBS and then returned to their original culture medium. The signals from both oxidized C11 (488 nm/FITC, laser/filter) and non-oxidized C11 (561 nm/Cy3, laser/filter) were monitored. Maximum projection images were generated. The ratio of the mean fluorescence intensity (MFI) of FITC to MFI of Cy3 was calculated as the relative LPO level by using Harmony software (Revvity, v5.1).

### Drug treatment

Stock solutions (50 mM) of resmetirom (MGL-3196, MedChem Express) were prepared from powder by resuspending in DMSO (Sigma Aldrich, 34869). The stock solution was first diluted in DMSO to make a 5 mM solution and then further diluted in perfusion media (EMS) to give the final concentration of 1 μM for drug treatment and 0.02% DMSO for the vehicle control. LAMPS were then maintained in EMS medium containing either vehicle control or the 1 μM resmetirom for 8 days at a flow rate of 5 μL/h. The images and collected efflux were analyzed and the drug treated devices were normalized to their respective vehicle control (described below). The drug binding capability of the polydimethylsiloxane (PDMS)-containing LAMPS device was assessed as previously described (46, 62, 63) using perfusion flow tests and mass spectrometry analysis of efflux collected from cell-free LAMPS devices after 72h of flow to determine the overall effective concentration of each compound compared to the starting concentration of drug in the perfusion medium.

### LipidTOX labeling and αSMA immunofluorescence

Cells were fixed with 4% paraformaldehyde (Thermo Fisher Scientific, AA433689M) in PBS for 30 min then washed twice with PBS for 10 min at room temperature. Following fixation, LipidTOX Deep Red Neutral Lipid Stain (1:500, Invitrogen, H34477) and mouse monoclonal anti-α-smooth muscle actin (αSMA) antibody (1:100, Sigma Aldrich, A2547) in PBS were perfused into devices and incubated overnight at 4°C. The following day, devices were washed twice with PBS and then incubated for 2 h with Alexa Fluor Goat anti-mouse 488 (1:250, Invitrogen, A-11029) secondary antibody and Hoechst (5 μg/mL, Invitrogen, H1399) at room temperature. Last, devices were washed three times with PBS before imaging. If not imaged immediately, the samples were stored at 4°C and imaged within one week.

### Confocal imaging and analysis

Confocal imaging was performed with the Phenix High Content Imaging platform (Revvity), using a 40X/0.75 hNA Air objective. A z-stacks of 70 μm distance (3 μm spacing between slices) were obtained across an array of 3 x 7 adjacent fields covering an area of 2.15 mm^2^ in the LAMPS device. Images for each condition were acquired using the same exposure time and laser power settings to ensure that intensity values were ∼50-90% of the total dynamic range. Image analysis was performed using custom analysis protocols developed in the Harmony (Revvity, v5.1) software package.

### Measurement of steatosis

Steatosis measurement was performed after completion of the experimental time course on Day 8. The nuclei were visualized by the staining with Hoechst and acquired using 405 nm laser and DAPI filter. The LipidTox signal was acquired using 640 nm laser and Cy5 filter. Imaging parameters (exposure time and laser power) were set using the EMS because this condition demonstrated the most LipidTOX labeling. LipidTox fluorescence intensity volume was calculated using 3D analysis method in Harmony software (Revvity, v5.1). Lipid droplet objects were identified using a local thresholding method and region scaling parameter defined by the Harmony software. This method creates a region or set of regions covering all pixels of the image with an intensity higher than their locally surrounding intensity.

### Measurement of stellate cell activation

Immunofluorescence for LX-2 cell expression of αSMA was performed after completion of the experimental time course on Day8. Nuclei signal was acquired using the 405 nm laser and DAPI filter. αSMA signal was acquired using 488 nm laser and FITC filter. Image analysis of LX-2 αSMA expression was quantified using a maximum intensity projection in the Harmony software (Revvity, v5.1). The particle detection function was then applied with a size exclusion setting of 100 μm^2^ to exclude non-specific labeling. The mean fluorescence intensity (MFI) of FITC was calculated as the integrated intensity of the αSMA signal.

### Efflux collection and biochemical measurements

Albumin (ALB), urea (BUN), lactate dehydrogenase (LDH), Pro-Collagen I Alpha 1 (COL1A1) were measured as previously described (46, 47, 62, 63). Briefly, efflux from LAMPS were collected on day 2, 4, 6, and 8. ALB assays were performed in a 1:100 efflux dilutions by enzyme linked immunosorbent assay (ELISA) using commercial antibodies (Bethyl Laboratories, A80-129A and A80-129P) and an ELISA accessory kit (Bethyl Laboratories, E101) with a human albumin standard prepared in house (MilliporeSigma, 126658). COL1A1 was measured using the Human pro-collagen 1A1 ELISA kit (R&D Systems, cat. no. DY6220-05) in a 1:50 efflux dilution. BUN was measured using the Stanbio BUN liquid reagent for diagnostic set (Stanbio Laboratory, cat. no. SB-0580-250). LDH was measured using the CytoTox 96 Non-Radioactive Cytotoxicity Assay (Promega, cat. no. G1780). The protocols for BUN and LDH assay were modified to a 384 well microplate format with no efflux dilution.

### Multiplex immunoassays

The levels of IL-6, IL-8, and MCP-1 were determined in efflux collected on day 8 using a custom version of the Human XL Cytokine Performance Panel (R&D systems). SHBG levels were quantified using the Human XL Cytokine Discovery Panel (R&D systems). Assays were completed according to the manufacturer’s instructions at The University of Pittsburgh Cancer Proteomics Facility Luminex® Core Laboratory on the xMAP platform. All the cytokine target profiling experiments were performed from efflux obtained from n = 3 devices for each experimental condition.

### Statistical analysis

Inter-study reproducibility analysis was performed in EveAnalytics^TM^ (previously, BioSystics Analytics Platform and the MPS-Database) using the Pittsburgh Reproducibility Protocol (PReP) (69, 70). Data analyzing genotype-specific differences across media conditions and drug treatments were analyzed and plotted in R (version 4.3.1). Data were obtained with a minimum of n = 3 LAMPS for each patient cell lot for each of the 3 media conditions and drug treatments. Data are plotted as mean ± standard error of mean (SEM). Statistical significance was assessed by ANOVA with Tukey’s test for multiple comparisons with a = 0.05 was considered statistically significant.

## Results

### Increased steatosis is observed in PNPLA3 rs738409 GG variant LAMPS compared to CC wild type LAMPS consistent with the clinical characterization of this polymorphism

To initiate the precision medicine approach outlined in **Figure 1**, we implemented the use of specific lots of patient hepatocytes that were genotyped to identify either PNPLA3 CC wild type or the rs738409 GG PNPLA3 high-risk variant **(Table S2**). These genotyped primary hepatocytes were used in combination with primary liver sinusoidal endothelial cells (LSECs) and cell lines for hepatic stellate cells (LX-2 cell line) and Kupffer-like cells (THP-1 cell line) to construct the LAMPS (**Fig S1A**). The impact of the PNPLA3 rs738409 GG variant on MASLD severity and progression coupled with lifestyle effects in LAMPS was then monitored over an 8-day time course by modulating specific media components including glucose, insulin, free fatty acids and immune activators **(Table S1)** to recapitulate MASLD progression from the normal fasting (NF) through early metabolic syndrome (EMS; early stage MASLD) and late metabolic syndrome (LMS; late stage MASLD) stages using various clinically relevant metrics including steatosis, immune activation, stellate cell activation, and fibrosis **(Fig S1B)** (46, 47).

To evaluate genotype-specific differences in both model functionality and cytotoxicity in each media condition, PNPLA3 GG variant and CC wild type LAMPS were maintained in either NF, EMS, and LMS media and efflux samples were collected on days 2, 4, 6 and 8 to measure the secretion of albumin (ALB) and blood urea nitrogen (BUN) for LAMPS functionality and lactate dehydrogenase (LDH) for LAMPS cytotoxicity **(Figure S2)**. While ALB secretion was higher in EMS medium compared to NF and LMS media, consistent with our previous work (46, 47), no significant differences in ALB secretion were observed between PNPLA3 GG variant and CC wild type LAMPS in any media condition **(Fig S2A)**, demonstrating similar overall model functionality. However, a significant increase in BUN secretion was observed on days 4, 6 and 8 in EMS medium in PNPLA3 GG LAMPS compared to CC wild type LAMPS, suggesting an overall increase in protein catabolism in LAMPS constructed with the high-risk variant in this medium condition **(Fig S2B)**. In addition, a significant increase in LDH secretion was observed in both NF and EMS media on day 8, and on days 4 and 6 in LMS medium, suggesting an overall increase of cytotoxicity in PNPLA3 GG LAMPS, consistent with its characterization as a high-risk variant associated with increased risk for MASLD progression and liver damage **(Fig S2C)** (71–73).

As excess hepatic fat content is strongly associated with the PNPLA3 GG variant (23, 71), we next evaluated genotype-specific effects on hepatocellular steatosis using quantitative fluorescence imaging to quantify LipidTOX staining in PNPLA3 GG variant or CC wild type LAMPS that were maintained for 8 days in either NF, EMS, and LMS media **(Figures 2 and S3)**. Compared to CC wild type LAMPS, significant increases in steatosis were observed in PNPLA3 GG variant LAMPS in all three media conditions including NF medium, indicating that increased hepatic lipid content is associated with the GG variant in both the baseline normal fasting (NF) condition as well as in both early and late-stage MASLD (EMS and LMS) media conditions **(Fig 2A and 2B)**, consistent with the clinical characterization that this high-risk variant is associated with increased susceptibility to hepatic steatosis (23, 72). In addition, significant increases in steatosis were observed in both the EMS and LMS media conditions compared to NF medium for both PNPLA3 CC wild type and GG LAMPS **(Fig S3)**, demonstrating that the EMS and LMS media formulations recapitulate lipid accumulation associated with the progression of MASLD.

**Figure 2.**
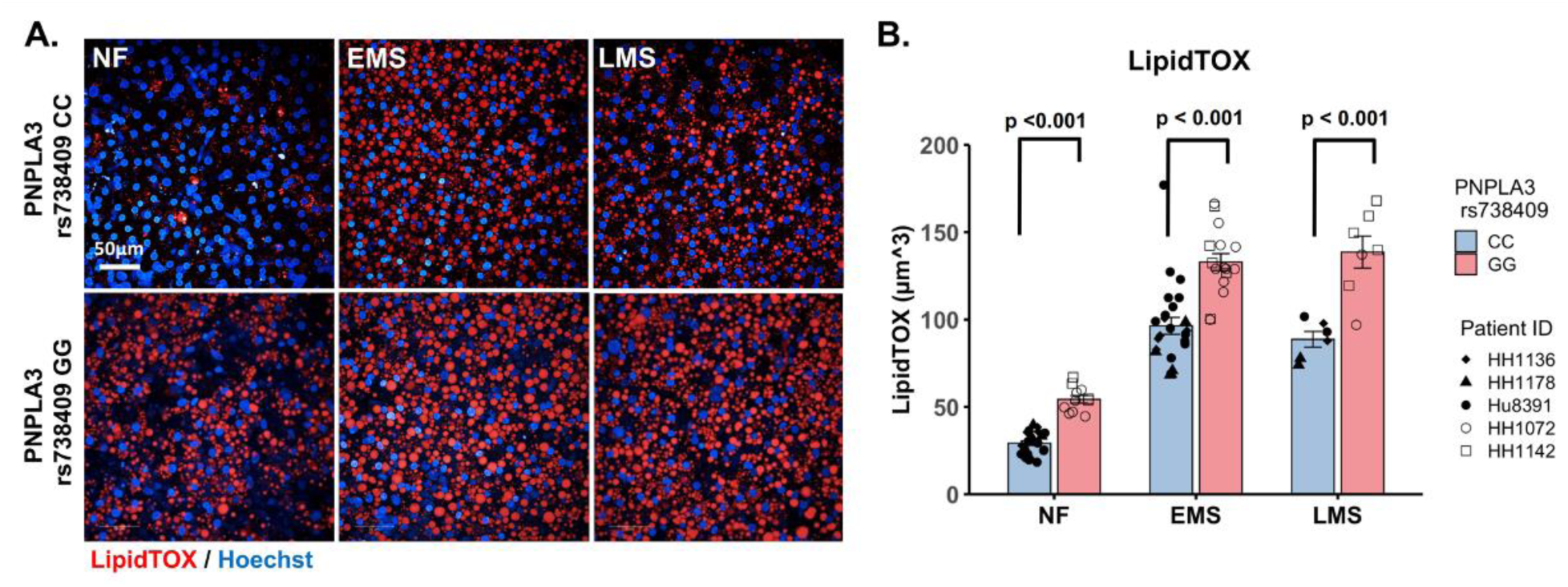
Increased steatosis in PNPLA3 rs738409 GG variant LAMPS compared to CC wild type LAMPS is consistent with the clinical characterization of the GG variant. (A) Representative images of LipidTOX labeled PNPLA3-LAMPS maintained in NF, EMS and LMS media. 40X; scale is 50µm. (B) Steatosis was quantified by quantitative fluorescence imaging of LipidTOX labeled samples in each media condition. Significant increases in steatosis were observed in PNPLA3 GG variant LAMPS compared to CC wild type LAMPS in all three media conditions, consistent with the clinical characterization of this high-risk variant (23). Data were obtained on Day 8 with a minimum of n = 3 LAMPS from each patient lot for each condition and plotted mean ± SEM. Statistical significance was assessed by ANOVA with Tukey’s test. p-value < 0.05 was considered statistically significant. Statistical comparisons across media conditions within each genotype were also performed in Figure S3.

Previous studies have shown that the increased steatosis sociated with MASLD progression impairs mitochondrial oxidative capacity resulting in the increased production of reactive oxygen species (ROS) (74, 75). Moreover, it has also recently been demonstrated that the PNPLA3 GG variant promotes the progression of MASLD by inducing mitochondrial dysfunction (76, 77). We next quantitatively assessed lipid peroxidation (LPO) in a 96-well plate format LAMPS (62) constructed with PNPLA3 GG variant or CC wild type hepatocytes and nonparenchymal cells (LSECs, LX-2 cells and THP-1 cells) that were maintained in NF, EMS, or LMS media for 5 days **(Fig S4)**. At the conclusion of the experimental time course, 96-well LAMPS were labeled with BODIPY 581/591 C11, a ratiometric fluorescent sensor for LPO, to determine the ratio of oxidized BODIPY (green) to reduced BODIPY (yellow) reflecting the overall LPO status within the model. Consistent with recent studies (76, 77), a significant increase in the ratio of Oxidized BODIPY/reduced BODIPY was observed in PNPLA3 GG variant 96-well plate LAMPS compared to the CC wild type in each media condition, demonstrating increased ROS production associated with the PNPLA3 GG variant **(Fig S4A and S4B)**.

### Increased production of pro-inflammatory cytokines is observed in PNPLA3 rs738409 GG variant LAMPS compared to CC wild type LAMPS demonstrating enhanced immune activation

The production of inflammatory cytokines is a significant factor contributing to the progression of MASLD (78, 79). Several experimental models demonstrate that the PNPLA3 rs738409 GG variant is associated with both elevated pro-inflammatory cytokine production as well as increased incidence of advanced fibrosis (44, 80, 81). To investigate the effect of the PNPLA3 rs738409 GG variant on immune activation, PNPLA3 GG variant and CC wild type LAMPS were maintained for 8 days in either NF, EMS, and LMS media and day 8 efflux samples were analyzed to measure the secretion of a panel of cytokines associated with MASLD progression, including CCL2, IL-6, and IL-8 (82, 83) **(Figures 3 and S5)**. Comparing absolute cytokine values between PNPLA3 LAMPS in each media type, no significant differences in CCL2 secretion were observed between PNPLA3 GG variant compared to CC wild type LAMPS in any media condition **(Fig 3A)**. A significant increase in IL-6 secretion was observed in the PNPLA3 GG variant compared to CC wild type in LMS medium (**Fig 3B**), and a significant increase in IL-8 secretion was observed in the PNPLA3 GG variant compared to CC wild type LAMPS maintained in NF and LMS media **(Fig 3C)**. The increases observed in LMS medium reflects a later stage of disease progression.

**Figure 3.**
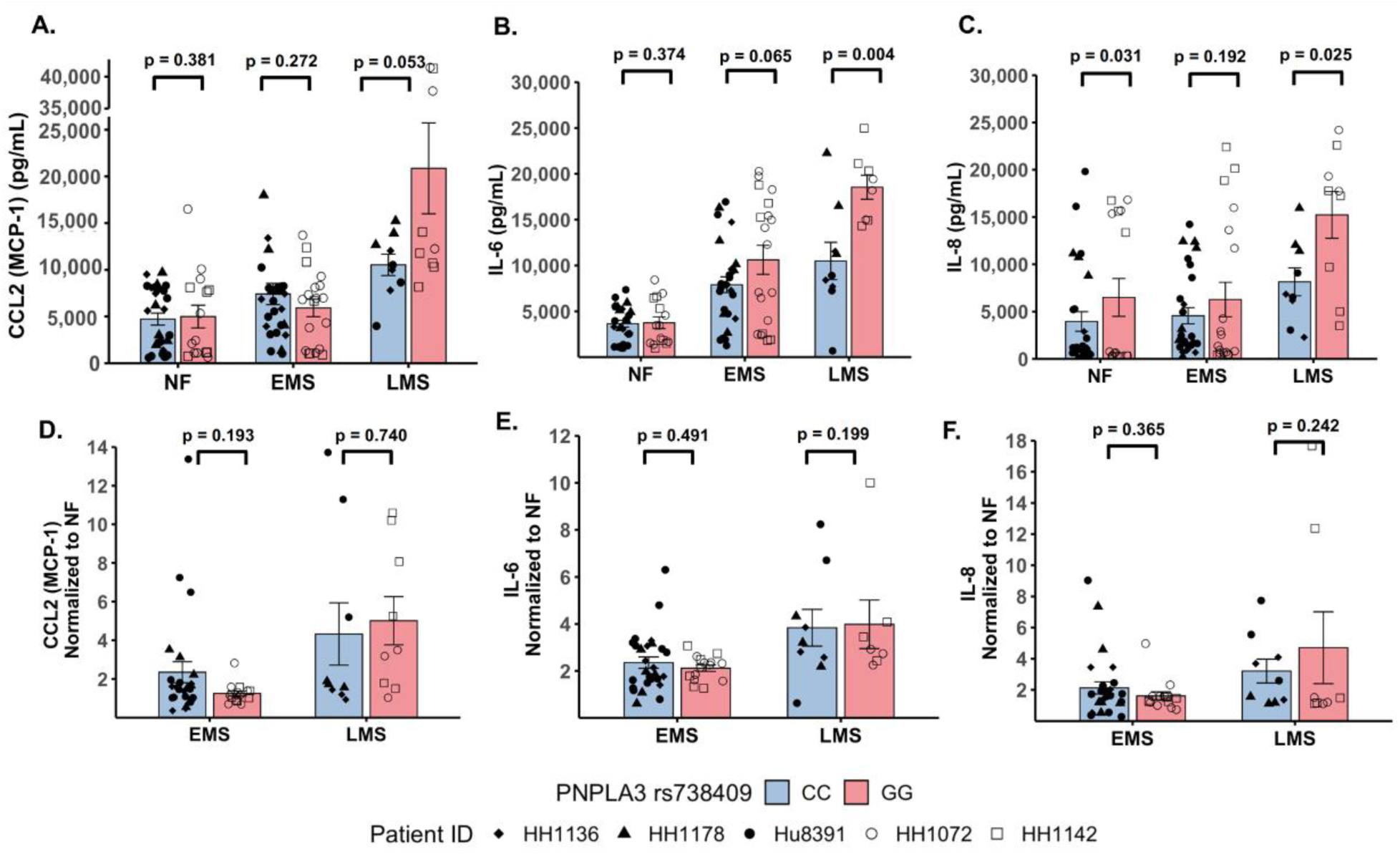
Increased pro-inflammatory cytokine production is observed in PNPLA3 rs738409 GG variant compared to CC wild type LAMPS, demonstrating enhanced immune activation. (A-C; absolute cytokine values) For NF medium, a significant increase in IL-8 secretion (C) was observed in PNPLA3 GG variant compared to CC wild type LAMPS. For LMS medium, significant increases in the secretion of both IL-6 (B) and IL-8 (C) were observed in PNPLA3 GG variant compared to CC wild type LAMPS, consistent with studies demonstrating increased immune activation and inflammation associated with the PNPLA3 GG variant (80, 81, 94). No significant differences in CCL2 secretion were observed between PNPLA3 GG variant compared to CC wild type LAMPS in any media condition. (D-F; normalized to NF values) Absolute cytokine values shown in panels A-C were normalized to their respective NF control for each study. For normalized cytokine values, while no significant differences between PNPLA3 GG and CC LAMPS were observed in either EMS and LMS media, the overall secretion levels of CCL2, IL-6, and IL-8 were all increased in EMS and LMS media compared to NF medium. The statistical comparisons between media types within each PNPLA3 genotype are shown in Figure S5. Data were obtained on Day 8 with a minimum of n = 3 LAMPS from each patient ID for each medium condition. Plotted is the mean ± SEM. Statistical significance was assessed by ANOVA with Tukey’s test. p-value < 0.05 was considered statistically significant.

In addition to the absolute cytokine values shown in **Figure 3A-C**, individual cytokine values were also normalized to their respective NF medium control from each individual study to determine the relative increases in individual cytokine production in EMS and LMS media **(Fig 3D-F)**. Though no significant differences between PNPLA3 GG and CC LAMPS were observed in either EMS and LMS media, the overall secretion levels of CCL2 **(Fig 3D)**, IL-6 **(Fig 3E)**, and IL-8 **(Fig 3F)** were all increased in EMS and LMS media compared to NF medium. We also performed statistical comparisons between media types within each PNPLA3 genotype and showed that the secretion of all three cytokines was significantly higher in LMS compared to NF media in both PNPLA3 CC wild type and GG variant LAMPS **(Fig S5A-C).** However, while all three cytokines were significantly increased in LMS compared to EMS media in PNPLA3 GG variant LAMPS, only IL-8 secretion was significantly increased in PNPLA3 CC wild type LAMPS in LMS medium **(Fig S5A-C)**. Taken together, these results demonstrate that the EMS and LMS media formulations recapitulate the immune activation associated with the progression of MASLD and increased pro-inflammatory cytokine production is observed in the PNPLA3 GG variant compared to CC wild type, consistent with the clinical characterization that the PNPLA3 rs738409 GG variant is associated with an increased risk of MASLD progression and advanced fibrosis (26, 81, 84–86).

### Increased stellate cell activation and COL1A1 secretion is observed in PNPLA3 rs738409 GG variant LAMPS compared to CC wild type LAMPS and is consistent with the increased incidence of advanced fibrosis in patients carrying the PNPLA3 high-risk allele

The PNPLA3 rs738409 GG variant is associated with increased risk for MASLD progression and advanced fibrosis (26, 81, 84, 85). We examined whether there were genotype-specific changes in stellate cell activation using fluorescence imaging to quantify the expression of αSMA, a marker for stellate cell activation, and production of the pro-fibrotic marker pro-collagen 1A1 (COL1A1) in PNPLA3 GG variant or CC wild type LAMPS that were maintained for 8 days in either NF, EMS, and LMS media **(Figures 4 and S6)**. For both analyses, an overall increase in αSMA fluorescence intensity **(Fig 4A and 4B)** and secretion of COL1A1 **(Fig 4C)** were observed in the PNPLA3 GG variant LAMPS compared to CC wild type LAMPS in all three media types with significant increases observed in both NF and LMS media. In addition, comparisons across media conditions within each PNPLA3 genotype demonstrated significant increases in αSMA fluorescence intensity in both EMS and LMS media conditions compared to NF medium for both PNPLA3 CC wild type and GG LAMPS **(Fig S6A)**. While PNPLA3 CC wild type LAMPS displayed a significant increase in COL1A1 secretion in both EMS and LMS media, COL1A1 secretion was significant only in LMS medium for PNPLA3 GG LAMPS **(Fig S6B)**. Taken together, these results demonstrate that the presence of the GG variant results in an elevated pro-fibrotic state in both normal fasting conditions as well as under MASLD conditions, consistent with the role of this high-risk variant being associated with an increased risk for progression to advanced fibrosis (26, 81, 84, 85).

**Figure 4.**
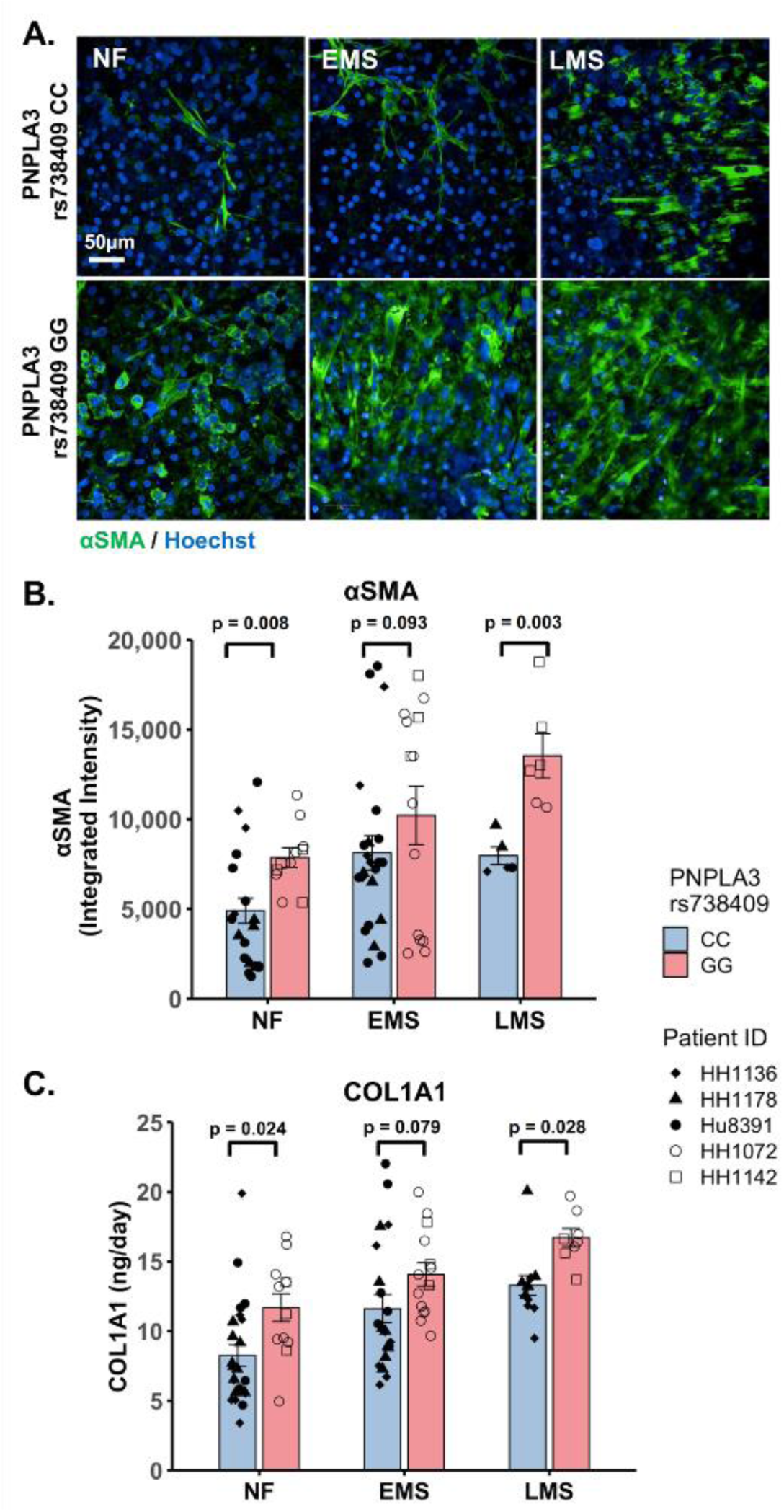
Increased stellate cell activation and COL1A1 secretion is observed in PNPLA3 GG variant LAMPS compared to CC wild type LAMPS demonstrating a more severe pro-fibrotic phenotype, consistent with the clinical characterization of the GG variant. (A) Representative images of αSMA labeled PNPLA3-LAMPS maintained in NF, EMS and LMS media. 40X; scale 50µm. (B) A Significant increase in αSMA integrated intensity was observed in PNPLA3 GG variant LAMPS compared to CC wild type LAMPS in both NF and LMS media, indicating an increase in stellate cell activation. (C) In addition, a significant increase in COL1A1 secretion was also observed in PNPLA3 GG variant compared to CC wild type LAMPS in both NF and LMS media. Taken together, these results demonstrate that the PNPLA3 variant is associated with increased stellate cell activation and subsequent secretion of pro-fibrotic markers (COL1A1), consistent with the clinical characterization of this high-risk variant (26, 81, 84, 85). Data were obtained on Day 8 with a minimum of n = 3 LAMPS from each patient lot for each condition and plotted mean ± SEM. Statistical significance was assessed by ANOVA with Tukey’s test. p-values < 0.05 were considered statistically significant. Statistical analysis comparing across media types for each PNPLA3 genotype were also performed and are shown in Figure S6.

### Both PNPLA3 rs738409 GG variant LAMPS and CC wild type LAMPS demonstrate overall excellent reproducibility for steatosis, immune activation, and fibrosis metrics in NF and EMS media when disease state, genotype and patient cohort are segmented

Reproducibility was assessed for both PNPLA3 GG and CC LAMPS for LipidTOX (steatosis), normalized cytokine values (immune activation), and α-SMA integrated intensity (stellate cell activation) for each patient cell lot used in the studies to generate **Figures 1-4**. The Numa Biosciences EveAnalytics^TM^ platform was implemented to determine the reproducibility score for each metric for Day 8 values. **Table 1** demonstrates that excellent reproducibility was observed in both NF and EMS media except for CCL2 secretion in patient lot 8391 among the PNPLA3 CC wild type patient cell lots as well as α-SMA integrated intensity in patient lot 1072 and IL-6 secretion in patient lot 1142 among the PNPLA3 GG variant patient cell lots. Overall, this analysis demonstrates that PNPLA3 LAMPS for both PNPLA3 genotypes are reproducible using a variety of MASLD-specific metrics in both NF and EMS media while only one individual study was performed for LMS medium using these patient cell lots, so reproducibility could not be assessed.

**Table 1.** PNPLA3-LAMPS demonstrate overall excellent reproducibility for steatosis, fibrosis, and immune activation when disease state, genotype and patient cohort are segmented. Reproducibility was assessed for αSMA integrated intensity, LipidTOX and normalized cytokine values for each patient lot used in the studies to generate Figures 1-4. Excellent reproducibility was observed for all media conditions except for αSMA integrated intensity in patient lot 1072, CCL2 secretion in patient lot 8391, and IL-6 secretion in patient lot 1142. The reproducibility score is determined using an ANOVA p-value on the Day 8 measurement where a p-value greater than 0.05 indicates that the means of the data points are similar (reproducible). The Reproducibility Status Scale displays p-value ranges from high (p-value > 0.1) to low (p-value < 0.05). Metadata and experimental data were captured in Numa Biosciences EveAnalytics^TM^ platform (70).

| Media | Metric | Unit | PNPLA3 CC patient lots<br>(wild type) |  |  | PNPLA3 GG patient lots<br>(high risk) |  |
| --- | --- | --- | --- | --- | --- | --- | --- |
|  |  |  | 1136 | 1178 | 8391 | 1072 | 1142 |
| NF | αSMA | Integrated Intensity | 0.24 | 0.20 | 0.73 | 0.02 | 0.29 |
|  | LipidTOX | μm<sup>3</sup> | 0.24 | 0.32 | 0.24 | 0.12 | 0.13 |
| EMS | αSMA | Integrated Intensity | 0.34 | 0.68 | 0.14 | 0.23 | 0.35 |
|  | LipidTOX | μm<sup>3</sup> | 0.22 | 0.96 | 0.26 | 0.33 | 0.62 |
| EMS | CCL2 | Normalized to NF | 0.14 | 0.54 | 0.01 | 0.22 | 0.27 |
|  | IL-6 | Normalized to NF | 0.89 | 0.39 | 0.08 | 0.55 | 0.00 |
|  | IL-8 | Normalized to NF | 0.51 | 0.21 | 0.60 | 0.11 | 0.47 |

### Resmetirom treatment resulted in a greater reduction of steatosis in PNPLA3 CC wild type LAMPS compared to GG variant LAMPS demonstrating genotype-specific inhibition of MASLD progression

Overall, the role of the PNPLA3 rs738409 variant in the response MASLD treatment remains uncertain (36). Using the same approach outlined in **Figure S1** to examine the effect of the PNPLA3 GG variant on MASLD disease progression, we next used both PNPLA3 GG variant and CC wild type LAMPS to evaluate the efficacy of resmetirom (Rezdiffra^TM^), a thyroid hormone receptor beta (THR-β) agonist recently approved for the treatment of MASLD (18, 87, 88). We determined that no detectable amount of resmetirom was adsorbed by the PDMS component of the LAMPS device **(Table S3)**. PNPLA3 GG variant or CC wild type LAMPS were maintained for 8 days in EMS media containing resmetirom (1μM) or vehicle control **(Fig S1C)**. The 1 µM concentration chosen for these studies is the approximate reported Cmax for Resmetirom (0.9 uM) (89). The effect of resmetirom on disease progression was evaluated using a panel of metrics including steatosis, immune activation, stellate cell activation, and fibrosis.

To evaluate LAMPS functionality and cytotoxicity under conditions of resmetirom treatment, efflux samples were collected on days 2, 4, 6 and 8 from PNPLA3 GG variant and CC wild type LAMPS to measure the secretion of albumin (ALB) and blood urea nitrogen (BUN) for LAMPS functionality and lactate dehydrogenase (LDH) for LAMPS cytotoxicity **(Figure S7)**. In both PNPLA3 GG variant and CC wild type LAMPS no significant changes were observed between resmetirom treatment and vehicle control for the secretion of ALB **(Fig S7A)**, BUN **(Fig S7B)**, or LDH **(Fig S7C)**, indicating that resmetirom does not adversely affect model functionality or cytotoxicity (18, 88).

We evaluated the on-target pharmacodynamic activity of resmetirom by quantifying the secretion of SHBG in PNPLA3 GG variant and CC wild type LAMPS treated with 1µM resmetirom as the sex hormone binding globulin (SHBG) is a direct transcriptional target of THR-β activity in the liver (88, 90). On day 8, resmetirom treatment resulted in significantly increased SHBG secretion in both GG variant and CC wild type LAMPS **(Fig S8A)**, supporting the on-target pharmacodynamic activity of resmetirom. In addition, PNPLA3 CC wild type LAMPS displayed significantly higher SHBG secretion compared to GG variant LAMPS **(Fig 5A).** We then evaluated genotype-specific effects on steatosis using fluorescence imaging to quantify LipidTOX staining in PNPLA3 GG variant or CC wild type LAMPS that were maintained for 8 days in EMS containing resmetirom (1μM) or vehicle control. While treatment with resmetirom significantly reduced steatosis in both PNPLA3 GG variant CC wild type LAMPS compared to their respective vehicle controls **(Fig S8B)**, a significantly greater reduction in steatosis was observed in PNPLA3 CC wild type LAMPS (∼33%) compared to PNPLA3 GG variant (∼20%) **(Fig 5B and 5C)**, indicating that resmetirom has better efficacy in PNPLA3 CC wild type LAMPS.

**Figure 5.**
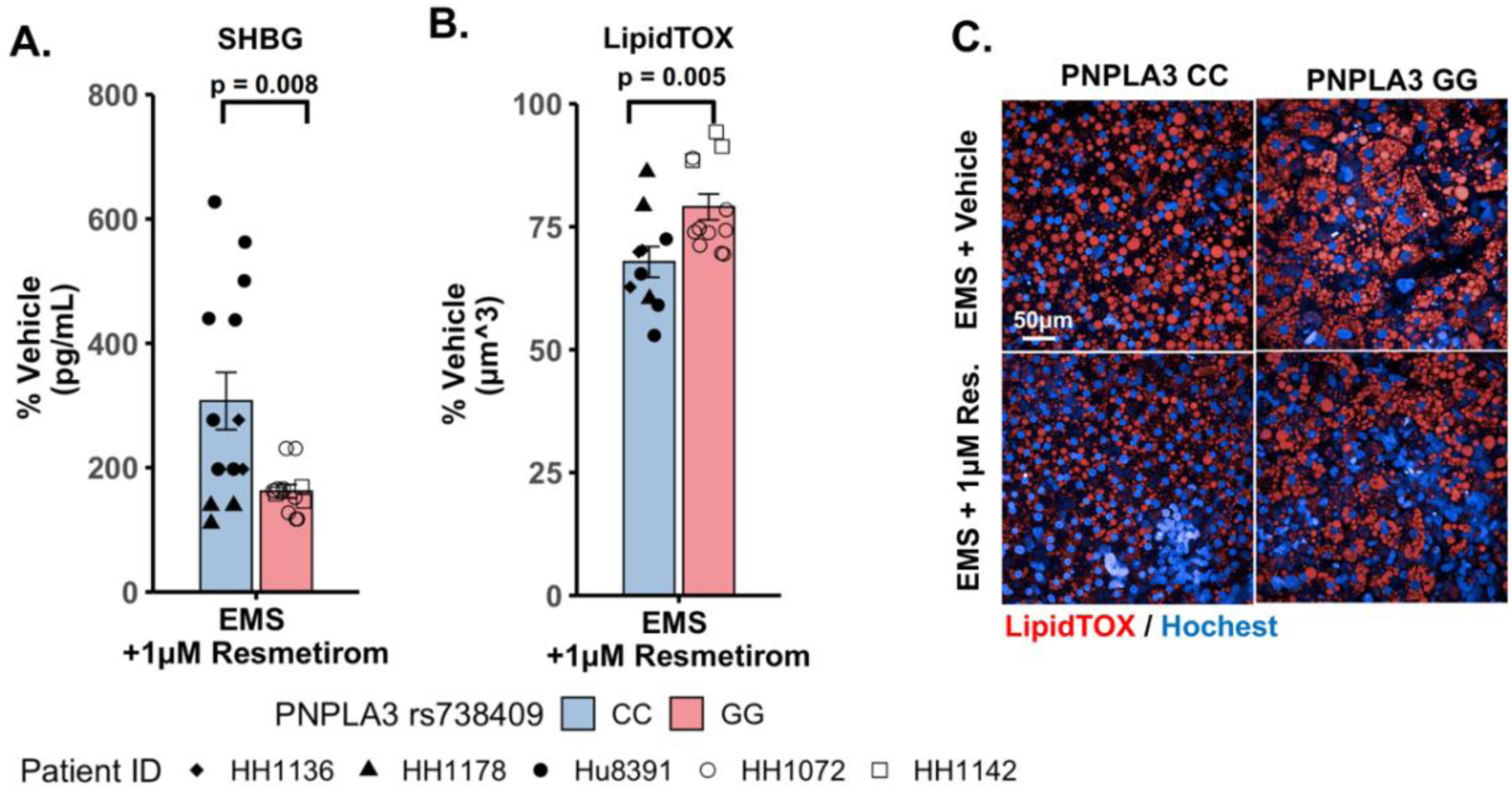
Resmetirom treatment resulted in a greater reduction of steatosis in PNPLA3 CC wild type LAMPS compared to GG variant LAMPS demonstrating genotype-specific inhibition of MASLD progression. PNPLA3 LAMPS models were maintained for 8 days in EMS media containing resmetirom (1μM) or DMSO vehicle control. The secretion of sex hormone binding globulin (SHBG) was assessed to determine the on-target pharmacodynamic activity of resmetirom and LipidTOX staining was quantified to determine the effect of resmetirom on steatosis. (A) 1 µM resmetirom treatment increased SHBG secretion in both PNPLA3-LAMPS, demonstrating the on-target pharmacodynamic activity of resmetirom. However, PNPLA3 CC wild type LAMPS displayed significantly higher SHBG secretion compared to GG variant LAMPS. (B) Resmetirom treatment resulted in a significantly greater reduction in steatosis in PNPLA3 CC wild type LAMPS (∼33%) compared to PNPLA3 GG variant (∼20%). (C) Representative images of LipidTOX labeled PNPLA3-LAMPS maintained in EMS with and without 1 µM resmetirom. 40X; scale 50µm. Data were obtained on Day 8 with n = 3 LAMPS from each patient lot for each condition and were plotted as average %vehicle ± SEM. Statistical significance was assessed by ANOVA with Tukey’s test. p-values <0.05 were considered statistically significant. Statistical analysis comparing resmetirom treatment with vehicle control within each PNPLA3 genotype was performed in Figure S8.

### Resmetirom treatment significantly reduced the secretion of the pro-inflammatory cytokine IL-6 in PNPLA3 GG variant LAMPS compared to CC wild type LAMPS

To evaluate potential differences in cytokine secretion in response to resmetirom treatment, PNPLA3 GG variant and CC wild type LAMPS were maintained for 8 days in EMS media containing resmetirom (1μM) or vehicle control and day 8 efflux samples were analyzed to measure the secretion of IL-6, CCL2, and IL-8. When the reduction of individual secreted cytokines (% of vehicle control) were compared directly between PNPLA3 GG variant and CC wild type LAMPS, no genotype-specific differences were observed **(Fig 6A-C)**. However, resmetirom treatment significantly reduced IL-6 secretion compared to vehicle control in PNPLA3 GG variant LAMPS (∼30% reduction), but not in PNPLA3 CC wild type LAMPS (∼15% reduction) **(Fig S9 B)**, while no reduction in either CCL2 **(Fig S9A)** or IL-8 **(Fig S9C)** was observed for either PNPLA3 LAMPS genotype relative to vehicle control. These results suggest that resmetirom has a larger effect on IL-6 secretion in the PNPLA3 GG variant.

**Figure 6.**
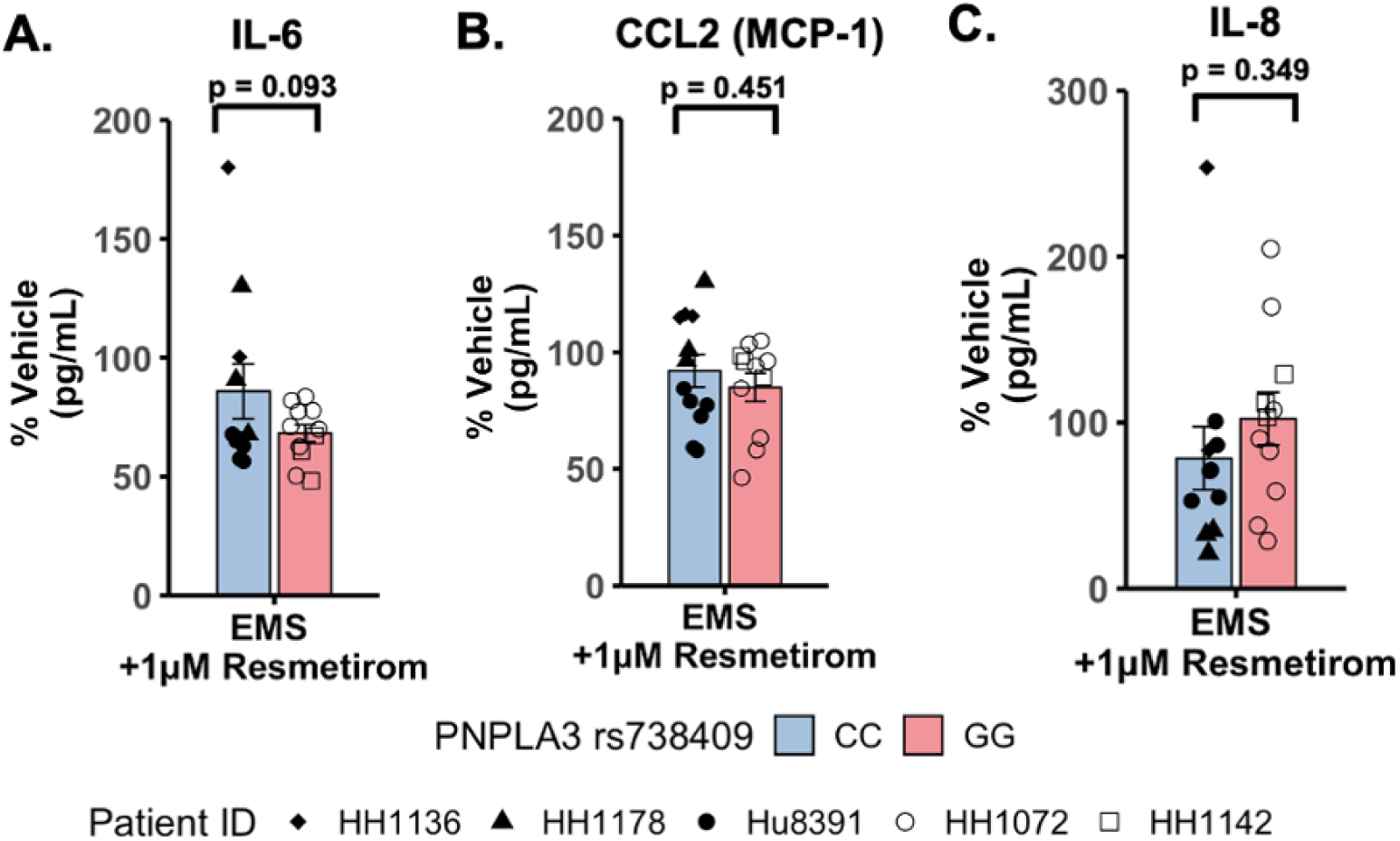
Resmetirom treatment did not result in any significant changes in pro-inflammatory reduction between PNPLA3 GG variant and CC wild type LAMPS. No significant differences in the reduction (% of vehicle control) of IL-6 (A), CCL2 (B), or IL-8 (C) were observed between PNPLA3 GG variant and CC wild type LAMPS upon treatment with 1µM resmetirom. Data (% of vehicle control) were obtained on Day 8 with a minimum of n = 3 LAMPS from each patient lot for each condition and plotted ± SEM. Statistical significance was assessed by ANOVA with Tukey’s test. p-values <0.05 were considered statistically significant.

### Resmetirom treatment resulted in a greater reduction of stellate cell activation and COL1A1 secretion in PNPLA3 CC wild type LAMPS compared to GG variant LAMPS demonstrating genotype-specific inhibition of MASLD progression

To evaluate genotype-specific differences in resmetirom treatment on the progression of pro-fibrotic phenotypes in PNPLA3 GG variant and CC wild type LAMPS, we quantified stellate cell activation (αSMA expression) using fluorescence imaging microscopy and pro-fibrotic marker secretion (COL1A1) in PNPLA3 LAMPS maintained for 8 days in EMS media containing resmetirom (1μM) or vehicle control. Resmetirom significantly reduced both αSMA integrated intensity **(Fig S10A)** and the secretion of COL1A1 **(Fig S10B)** compared to vehicle control in PNPLA3 CC wild type LAMPS, but not GG variant LAMPS. In addition, a significantly greater reduction in both αSMA integrated intensity (50% vs. 25%) **(Fig 7A and B)** and the secretion of COL1A1 (35% vs. 0%) **(Fig 7C)** was observed in PNPLA3 CC wild type LAMPS when compared to the PNPLA3 GG variant, demonstrating better resmetirom efficacy for both stellate cell activation and production of pro-fibrotic markers.

**Figure 7.**
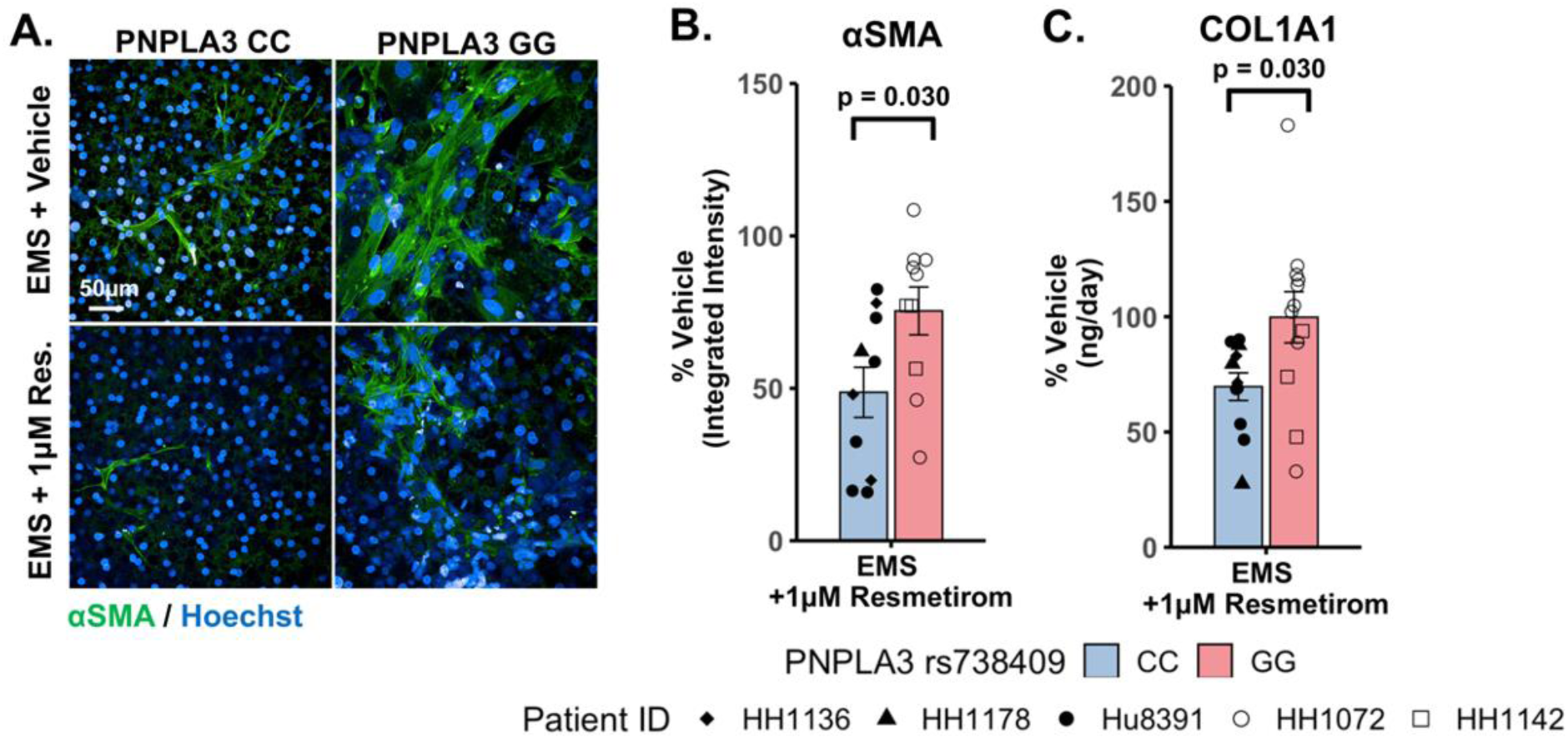
Resmetirom treatment resulted in a greater reduction of stellate cell activation and COL1A1 secretion in PNPLA3 CC wild type LAMPS compared to GG variant LAMPS. (A) Representative images of αSMA labeled PNPLA3-LAMPS maintained in EMS with 1µM resmetirom or vehicle control. 40X; scale 50µm. (B and C) A significantly greater reduction in both αSMA integrated intensity (B; 50% vs. 25%) and the secretion of COL1A1 (C; 35% vs. 0%) was observed in PNPLA3 CC wild type LAMPS compared to PNPLA3 GG variant LAMPS. Data were obtained on Day 8 with n = 3 LAMPS from each patient lot for each condition and were plotted as average %vehicle ± SEM. Statistical significance was assessed by ANOVA with Tukey’s test. p-values <0.05 were considered statistically significant. Statistical analysis comparing resmetirom treatment with vehicle control within each genotype was performed in Figure S10.

## Discussion

The clinical variability and pathophysiologic heterogeneity of MASLD has contributed to the challenges in therapeutic development of this disorder. To begin to address this challenge, we have initiated a MASLD precision medicine approach that harnesses the use of the LAMPS constructed with primary hepatocytes genotyped for the wild type PNPLA3 and the MASLD-associated genetic variant PNPLA3 rs738409. The initial goal was to establish the disease progression, severity, and response to drug treatment in a primary cell-focused model where we have fully adult hepatocytes and LSECs combined with well characterized human cell lines for stellate and Kupffer cells (40, 46, 47, 61–63). This model serves to develop the platform for evaluating the role of important gene variants, as well as lifestyle factors in MASLD disease progression and response to therapeutics. In addition, this model serves as a critical reference for evaluating the functionality and disease characteristics of our patient-derived, liver cell MPS (patient biomimetic twins or PBTs) (40, 46, 47, 75, 91, 92). We are beginning to implement the use of PBTs creating a precision medicine platform that will be used in characterizing patient cohorts for drug discovery, development (including clinical trials on chips) and precision therapeutic treatments **(Fig 1)**.

We show here that the LAMPS model recapitulates the pathophysiologic stages of early and late MASLD along with key differences in both disease progression and therapeutic responses in LAMPS constructed with either wild type or high-risk PNPLA3 alleles. To quantitatively interpret the data on disease progression and response to drug treatment the model had to be reproducible (67–70). The LAMPS demonstrated overall excellent reproducibility for steatosis, fibrosis, and immune activation when disease state, genotype and patient hepatocyte lot cohorts were segmented **(Table 1).** Our results demonstrating differences in PNPLA3 CC wild type compared to GG LAMPS are consistent with clinical studies in humans (20, 29, 81). The high-risk PNPLA3 GG variant LAMPS demonstrated increased steatosis, immune activation, stellate cell activation and secretion of the pro-fibrotic marker COL1A1 **(Table 2)**, consistent with the clinical characterization of this polymorphism. In addition, we observed genotype-specific differences in response to drug treatment, as resmetirom demonstrated greater efficacy in PNPLA3 wild type CC LAMPS compared to the GG variant in the reduction of steatosis, stellate cell activation and the secretion of marker COL1A1 (**Table 3)**.

**Table 2.** A summary of the MASLD progression in PNPLA3 CC wild type and GG variant LAMPS normalized to PNPLA3 CC wild type in NF.

| Metric | Condition | CC | GG | *P-value* |
| --- | --- | --- | --- | --- |
| Lipid Accumulation (μm<sup>3</sup>) | NF | 1.00 | 1.86 | <0.001 |
|  | EMS | 3.30 | 4.54 | <0.001 |
|  | LMS | 3.03 | 4.74 | <0.001 |
| IL-6 (pg/mL) | NF | 1.00 | 1.03 | 0.374 |
|  | EMS | 2.16 | 2.91 | 0.065 |
|  | LMS | 2.87 | 5.07 | 0.004 |
| IL-8 (pg/mL) | NF | 1.00 | 1.64 | 0.031 |
|  | EMS | 1.15 | 1.59 | 0.192 |
|  | LMS | 2.06 | 3.85 | 0.025 |
| CCL2 (pg/mL) | NF | 1.00 | 1.06 | 0.381 |
|  | EMS | 1.57 | 1.25 | 0.272 |
|  | LMS | 2.23 | 4.42 | 0.053 |
| αSMA (Integrated Intensity) | NF | 1.00 | 1.60 | 0.008 |
|  | EMS | 1.66 | 2.08 | 0.093 |
|  | LMS | 1.63 | 2.76 | 0.003 |
| COL1A1 (ng/day) | NF | 1.00 | 1.42 | 0.024 |
|  | EMS | 1.41 | 1.70 | 0.079 |
|  | LMS | 1.61 | 2.02 | 0.028 |

**Table 3.** A summary of genotype-specific response to resmetirom in PNPLA3 CC wild type and GG variant LAMPS.

|  | Resmetirom Treatment<br>in LAMPS<br>(% Reduction) |  | Comparison to Phase III Clinical<br>Trial (18) | Reduction with<br>resmetirom treatment to<br>vehicle control (%) |
| --- | --- | --- | --- | --- |
| Metric | CC | GG | General Population |  |
| Lipid Accumulation | 32.16 | 20.97 | ~ 30% reduction in liver fat |  |
| αSMA | 51.31 | 24.60 | ~ 25% patient response with<br>fibrosis improvement or no<br>worsening |  |
| COL1A1 | 30.34 | 0.23 |  |  |
| IL-6 | 14.08 | 31.71 | No available cytokine result |  |
| IL-8 | 21.49 | -2.27 |  |  |
| CCL2 | 7.98 | 15.05 |  |  |

The PNPLA3 polymorphism is associated with increased susceptibility to MASLD, disease progression to cirrhosis, and risk of developing HCC and prior research supports incorporation of PNPLA3 genotype into prognostic scores for predicting risk of disease progression (80, 81). Our experimental results demonstrate that the PNPLA3 rs738409 GG variant LAMPS exhibit increased steatosis in NF, EMS and LMS media compared to wild type CC LAMPS consistent with the clinical characterization describing abnormal lipid homeostasis in these patients (23, 71, 72, 76, 93). Interestingly, our results demonstrated increased pro-inflammatory cytokine production in PNPLA3 GG variant LAMPS, consistent with recent results obtained in both animal models and MPS (44, 80). However, a recent clinical study that examined serum cytokine levels in a cohort of 123 genotyped patients found no significant impact of the PNPLA3 polymorphism on cytokine levels within this cohort (94). Thus, additional larger clinical studies and further *in vitro* studies are needed to more fully determine whether cytokine profiles are impacted by the GG variant. Furthermore, our data also show that LAMPS constructed with the high-risk PNPLA3 polymorphism exhibit higher states of stellate cell activation and secretion of COL1A1, consistent with the association this variant with increased risk of advanced fibrosis (26, 84). Our findings have immediate implications for precision risk stratification, enrichment of patient cohorts for clinical trials, and selection of approved therapies for management of patient sub-groups with MASLD.

An overarching goal of our research is to develop a qualified human liver MPS drug discovery tool (DDT) to address several important contexts of use (CoU) including toxicology, drug discovery/profiling and clinical trials on chips. An immediate question is whether the MASLD LAMPS model could be used as a tool for segmenting patient cohorts based on response to approved drugs, drug candidates, (before or after entering the clinic, or failed due to missing endpoints in general patient populations), as well as predicted drugs for repurposing (40, 46). Our strategy has been to first focus on early stages of MASLD progression with stage 3 or less fibrosis that is maintained in the EMS media. Our first step was to determine if the PNPLA3 GG variant LAMPS and wild type CC LAMPS in the early metabolic syndrome (EMS) “lifestyle” state would respond distinctly to the recently approved drug, resmetirom, a liver-targeted thyroid hormone receptor (THR-β) selective agonist designed to target causes of MASH with moderate to advanced liver fibrosis (18, 87, 88). The initial study reported here was to determine the level of inhibition of disease progression over 8 days with a single dose (1 µM) based on the validated panel of metrics including steatosis, immune activation, and early fibrosis (stellate cell activation and secretion of COL1A1). We observed genotype-specific differences in response to resmetirom treatment, demonstrating that multiple MASLD disease progression metrics were significantly reduced in PNPLA3 wild type compared to GG LAMPS, including steatosis, stellate cell activation, and the secretion of a pro-fibrotic marker. Thus, these findings are consistent with the recent study where ∼15% of a general population in Phase III exhibited improvement in fibrosis or no worsening with resmetirom treatment (18).Our study has several strengths including that the use of genotyped primary hepatocytes that serve as an important benchmark for future studies using MPS constructed with iPSC-derived liver cells from patients (PBTs). A critical question in using iPSC-derived cells as a precision medicine tool has been the level of overall maturity and functionality and this requires a benchmark. Another strength of our study is that it demonstrates genotype-specific differences in both MASLD progression and response to drug treatment, indicating that a reproducible patient-derived MPS (PBT) could be harnessed to identify disease-state specific biomarkers that would optimize patient subgroups for therapeutic testing strategies.

One limitation of this study was that we tested two patient lots of PNPLA3 GG variant hepatocytes in LAMPS that were constructed with non-isogenic NPCs. Therefore, the key to creating a powerful precision medicine platform is the construction of patient biomimetic twins constructed with isogenic iPSCs derived from individual patients, a program that is in progress. Secondly, the addition of resmetirom to EMS medium in our present study occurred at the outset of establishing flow in the LAMPS to test the effect of resmetirom on preventing disease progression. Future studies using the existing hybrid LAMPS model, while we validate the PBTs, will evaluate the effect of resmetirom on reversing MASLD phenotypes in LAMPS where EMS disease features have been established. Additional steps will include full dose response curves on a larger genotyped pool of patient backgrounds including PNPLA3 and other known MASLD-associated variants. Individual drug candidates or combinations of drugs after the MASLD LAMPS have been progressed to EMS and LMS “lifestyle states” will be tested to explore if later stages of disease progression can be halted and/or reversed. Furthermore, the present model and the future PBTs will play a critical role in defining mechanisms of action of drug candidates and drugs.

Overall, there are many efforts toward developing precision medicine platforms for MASLD, including a variety of MPS designs, organoids, and humanized murine models. In addition, a combination of small molecules, antisense oligonucleotides, and siRNA-based silencing of other MASLD relevant genes and targets are under investigation (11, 95). Looking forward, we will continue the characterization of iPSC-derived liver cells from patients that have been enrolled in the University of Pittsburgh Medical Center Liver Steatosis and Metabolic Wellness Clinic and their incorporation into PBTs **(Fig. 1)** (40, 75, 92, 96). The reproducibility, functionality and response to disease progression and drug treatments will be benchmarked to the primary cell-focused study reported here.

We have patient clinomics data and are beginning to develop a variety of omics data on selected patients for clinical reference and future computational modeling (patient digital twins; PDTs). The goal is to produce cohorts of PBTs where the genotype, lifestyle, and environment histories, as well as co-morbidities are known. Having specific cohorts of patients and their PBTs representing some of the major sources of MASLD patient heterogeneity will facilitate the development of advanced qualified liver MPS drug development tools (DDTs) with many contexts of use (CoUs) that will aid drug discovery and development, including optimized clinical trials on chips by selection of high probability responders before testing in patients. Based on a growing set of clinomic and omic datasets on the enrolled patients, we will be in a position to extend our combined human MPS and quantitative systems pharmacology strategy (39, 97) to include the development of Patient Digital Twins (PDTs) to complement the PBTs for predicting patient cohort-specific toxicity and efficacy. We project that the integration of PDTs and PBTs for MASLD and other diseases will play critical roles in the advancement of precision medicine efforts.

## Author contributions

Conceptualization: D.L.T., M.T.M., A.S.G., J.B., and A.M.S.; data curation: M.X., M.V., I.P.P. and M.T.M.; formal analysis: M.X., M.V., I.P.P., M.S., and M.T.M.; funding acquisition: M.T.M., J.B., A.S.G. and D.L.T.; investigation: M.X., M.V., I.P.P., J.A.B., R.D., D.C.G., G.L., R.F., L.A.P.F. and M.S.; methodology: M.S., A.M.S., A.G. and M.T.M.; project administration: D.L.T., A.G., and C.R.; resources: C.R.; software: M.X., M.V., I.P.P. and D.C.G.; visualization: M.V., M.X., and D.C.G.; supervision: D.L.T., M.T.M., A.S.G., J.A.B., J.B., A.G., A.M.S. and M.S.; validation: M.S., L.A.V., A.M.S., A.G., A.S.G., and M.T.M.; writing-original draft: M.X., M.V., M.T.M and D.L.T.; writing-review and editing: I.P.P., D.C.G., R.D., G.L., R.F., L.A.P.F., C.R., M.S., A.M.S., J.A.B., L.A.V., A.G., J.B., A.S.G., M.T.M and D.L.T. All authors have read and agreed to the published version of the manuscript.

## Acknowledgements

The authors would like to thank all members of their laboratories for supporting the research efforts reported here. Gentest® 999Elite™ human hepatocytes were obtained from the Discovery Life Science. The cytokine data was obtained using the University of Pittsburgh Cancer Institute (UPCI) Cancer Biomarkers Facility: Luminex Core Laboratory. Mass spectrometry for PDMS drug binding studies was performed by Patrick Oberly at the University of Pittsburgh Small Molecule Biomarker Core (NIH grant S10OD028540: “Small Molecule Biomarker Core: TSQ Altis LC-MS/MS”). We acknowledge the use of the PLRC cores including the Human Synthetic Liver Biology Core. This project used shared instrumentation that was acquired with NIH grant S10OD028450. Figure 1 was created using bioRender.com.

