## Supplemental Material for "Comparison of Wild-Type and High-risk PNPLA3 variants in a Human Biomimetic Liver Microphysiology System for Metabolic Dysfunction-associated Steatotic Liver Disease Precision Therapy"

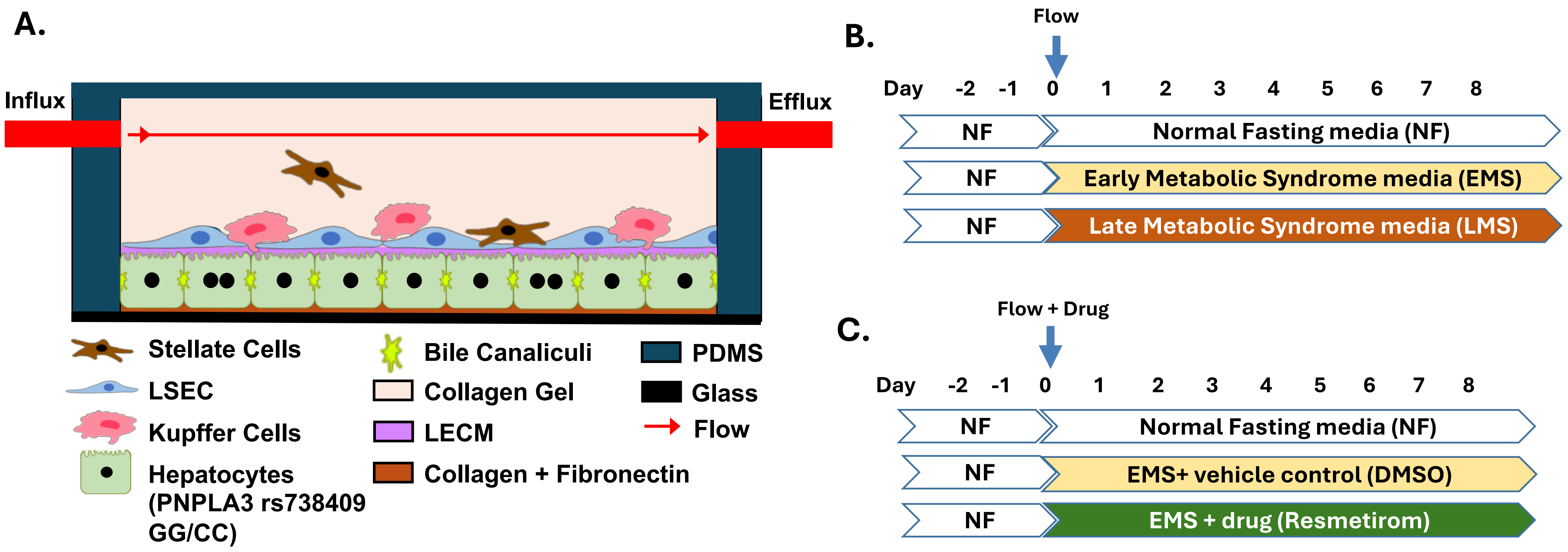


**Figure S1. Overview of the experimental setup for LAMPS MASLD disease progression and drug testing studies.** (A) LAMPS were constructed with 4 liver cell types sequentially layered starting with primary hepatocytes (PNPLA3 rs738409 GG/CC) followed by the addition of non-parenchymal cells (NPCs) including primary liver sinusoidal endothelial cells (LSECs), and LX-2 (hepatic stellate cells) and activated THP-1 (Kupffer-like cells) cell lines as previously described (1-5). (B-C) Experimental timeline used to monitor disease progression and drug testing in LAMPS using the previously described media formulations shown in Table S1 (NF, EMS, and LMS). For these studies, LAMPS were maintained at a flow rate of 5 μL/h over an 8-day time course to mimic zone 3 oxygen tension (2). For drug testing studies, resmetirom was added to EMS medium at time 0 and compared to a 0.02% DMSO vehicle control to evaluate the efficacy of this drug on MASLD progression.

**Table S1.** The formulations of NF, EMS and LMS media drive MASLD progression based on the effect of lifestyle on blood chemistry.

| Medium Component | NF (Normal Fasting) | EMS (Early Metabolic Syndrome; MASLD) | LMS (Late Metabolic Syndrome; MASH, T2D) |
|---|---|---|---|
| **Glucose** | 5.5 mM | 11.5 mM | 20 mM |
| **Insulin** | 10 pM | 10 nM | 10 nM |
| **Glucagon** | 100 pM | 10 pM | 10 pM |
| **Oleic acid** | - | 200 µM | 200 µM |
| **Palmitic acid** | - | 100 µM | 100 µM |
| **Lipopolysaccharide (LPS)** | - | - | 0.25 µg/mL |
| **Transforming Growth Factor β1 (TGF-β1)** | - | - | 5 ng/mL |

Media formulations were designed to mimic disease progression from the normal fasting (NF) to early metabolic syndrome (EMS; MASLD) and late metabolic syndrome (LMS; MASH, T2D) state. These media formulations were developed using glucose-free Williams E base medium supplemented with physiologically relevant levels of glucose, insulin, glucagon, oleic acid, palmitic acid and molecular drivers of fibrosis including TGF-β1 and LPS (4).

**Table S2:** Clinical characteristics of PNPLA3 rs738409 GG variant and CC wild type primary hepatocytes used in LAMPS studies.

| Vendor | Lot ID | Genotype (PNPLA3 rs738409) | Age | Sex | Race | BMI | Smoking | Alcohol | Infectious diseases* |
|---|---|---|---|---|---|---|---|---|---|
| **DLS** | HH1072 | GG | 40 | Female | Caucasian | 37.3 | Yes | Yes | HBV-, HCV-, HIV-, CMV- |
| **DLS** | HH1142 | GG | 27 | Female | Caucasian | 25 | No | No | HBV-, HCV-, HIV-, CMV+ |
| **DLS** | HH1178 | CC | 66 | Female | Caucasian | 31.6 | Yes | No | HBV-, HCV-, HIV-, CMV+ |
| **DLS** | HH1136 | CC | N/A | Male | Caucasian | 27.34 | N/A | N/A | N/A |
| **Thermo Fisher** | Hu8391 | CC | 62 | Male | Caucasian | 22.2 | Yes | Yes | HBV-, HCV-, HIV-, CMV- |

*Discovery Life Sciences, DLS; Hepatitis B Virus, HBV; Hepatitis C Virus, HCV; Human Immunodeficiency Virus, HIV; Cytomegalovirus, CMV.

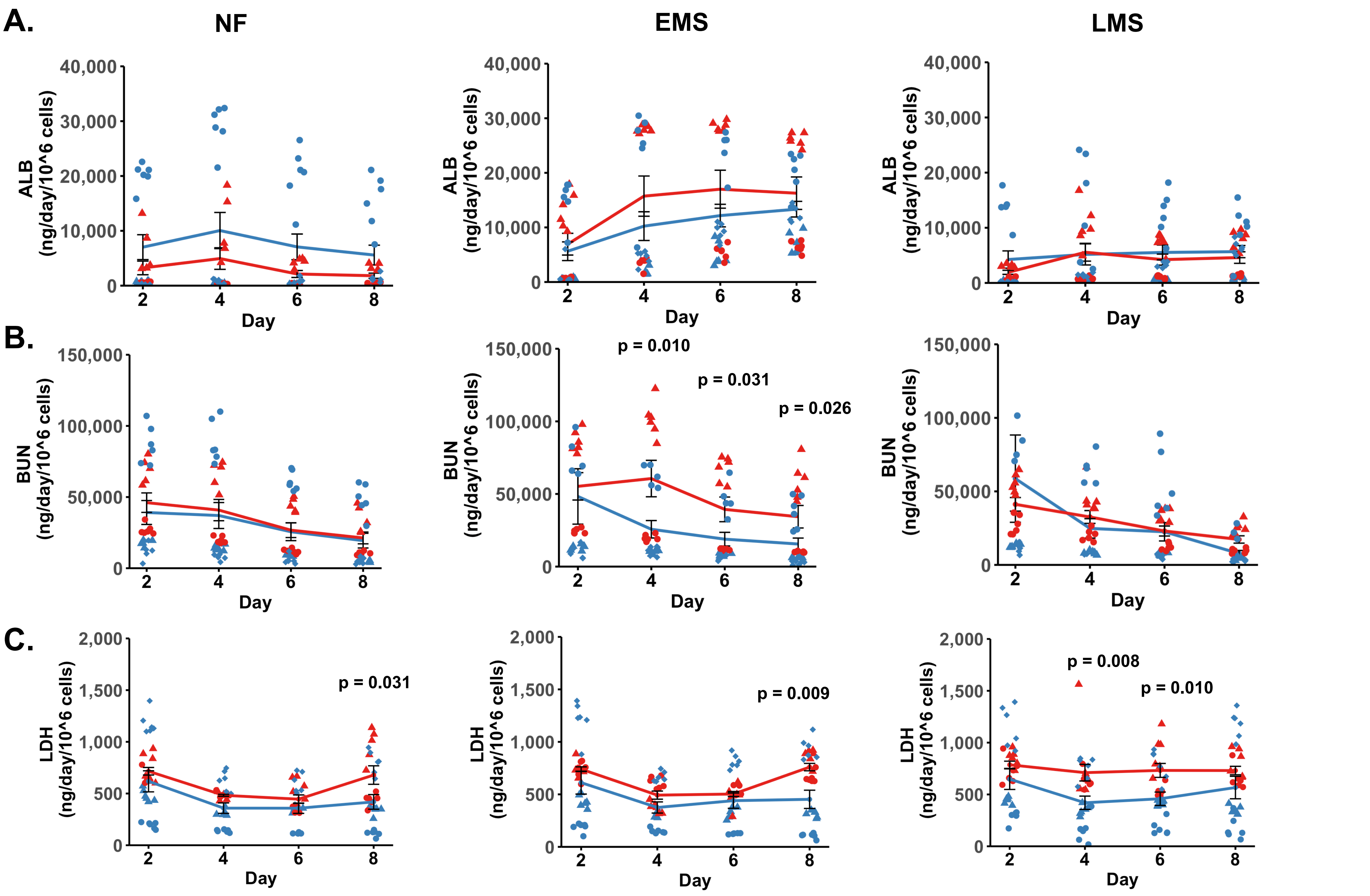


**Figure S2. While PNPLA3 rs738409 GG variant and CC wild type LAMPS exhibited similar overall model functionality, PNPLA3 GG LAMPS displayed increased cytotoxicity across media types.** (A-C) The secretion of albumin (ALB; A), blood urea nitrogen (BUN; B), and lactate dehydrogenase (LDH; C) were compared over the 8-day time course in PNPLA3 GG variant and CC wild type LAMPS to identify genotype-specific differences in LAMPS functionality (ALB and BUN) and cytotoxicity (LDH) in each media type (NF, EMS, LMS). While no significant changes in ALB secretion (A) were observed between PNPLA3 GG variant and CC wild type LAMPS, indicating similar overall model functionality, a significant increase in BUN secretion (B) was observed on days 4, 6 and 8 in EMS medium in PNPLA3 GG LAMPS. (C) A significant increase in LDH secretion was observed in both NF and EMS media on day 8, and on days 4 and 6 in LMS medium, suggesting an overall increase cytotoxicity in PNPLA3 GG LAMPS, consistent with its characterization as a high-risk variant. Data were plotted mean ± SEM from a minimum of n = 3 LAMPS from each patient lot for each media condition. Statistical significance was assessed by ANOVA with Tukey's test for each indicated time point. Only p-values < 0.05 were considered statistically significant and are indicated.

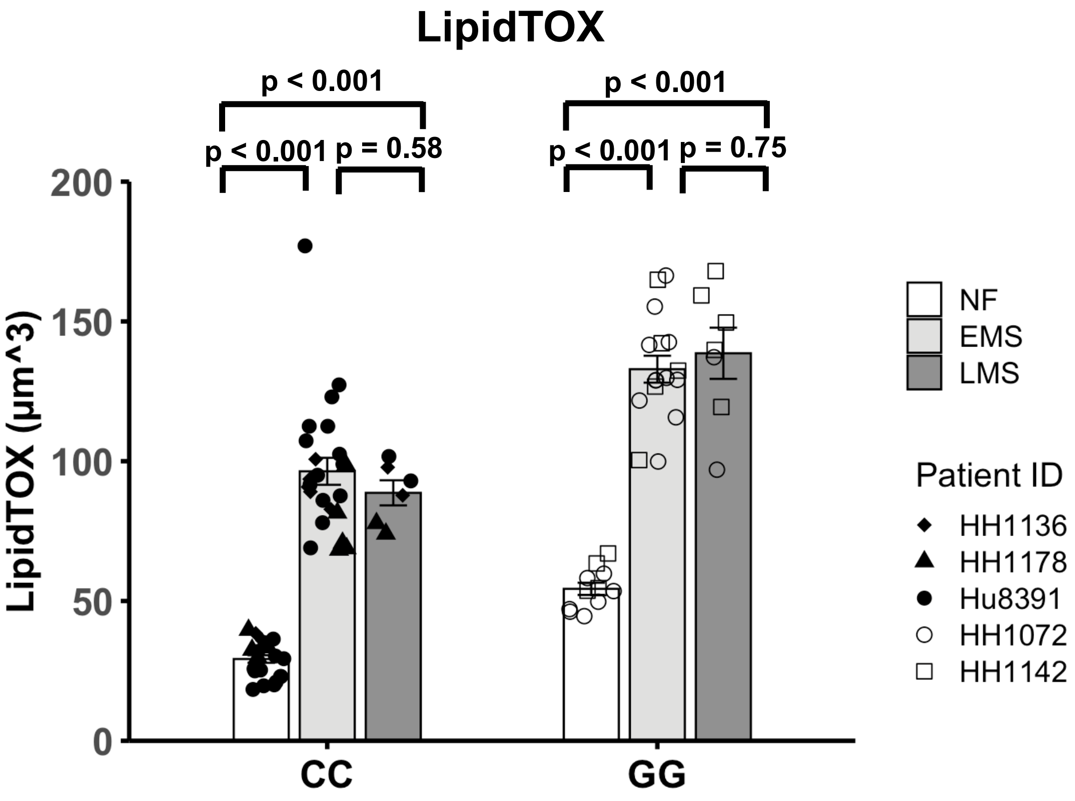


**Figure S3. Increased steatosis was observed in both EMS and LMS media compared to NF medium in both PNPLA3 CC and GG LAMPS, demonstrating lifestyle-induced MASLD progression.** Steatosis was quantified by quantitative fluorescence imaging of LipidTOX labeled samples in each media condition. Significant increases in steatosis were observed in both EMS and LMS media compared to NF medium within each PNPLA3 genotype. Data were obtained on Day 8 with a minimum of n = 3 LAMPS from each patient lot for each condition and plotted mean ± SEM. Statistical significance was assessed by ANOVA with Tukey's test. p-value < 0.05 was considered statistically significant. These data support the analysis performed in Figure 2.

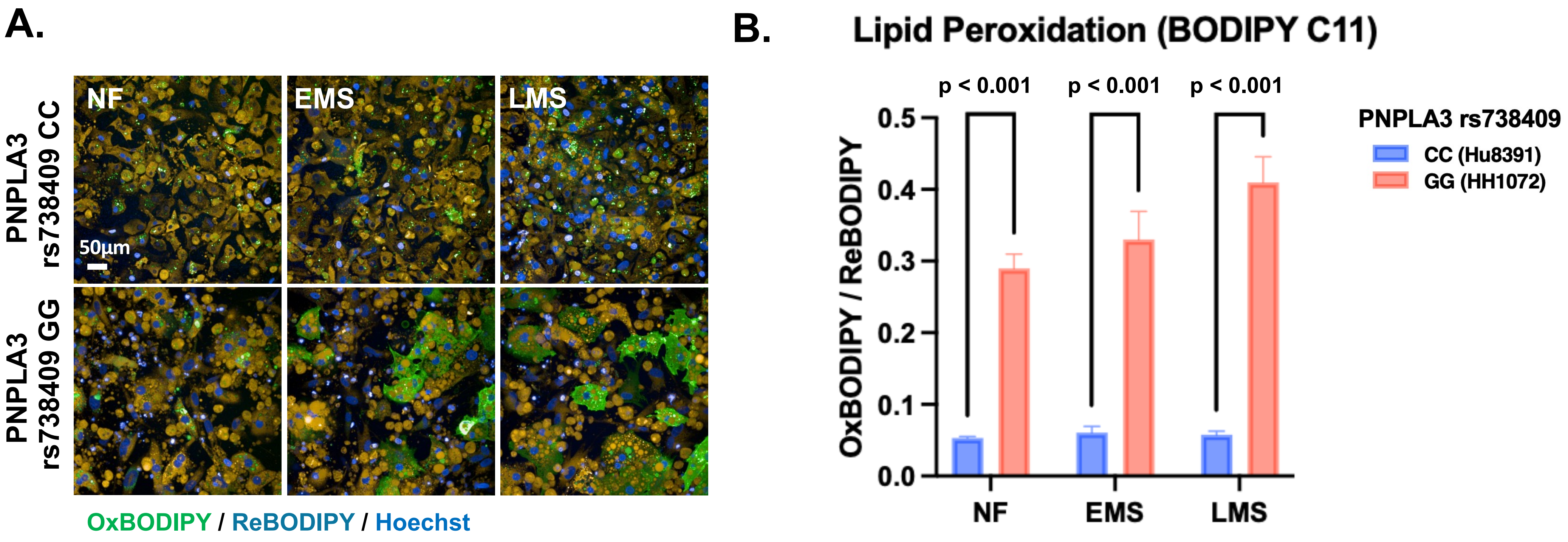


**Fig S4. Increased lipid peroxidation (LPO) was observed in PNPLA3 rs738409 GG variant 96-well plate LAMPS compared to PNPLA3 CC wild type demonstrating the role of oxidative stress in genotype-specific MASLD progression.** 96-well plate LAMPS were constructed using PNPLA3-genotyped hepatocytes and NPCs in NF, EMS and LMS media (6). Cells were labeled with BODIPY 581/591 C11 to monitor LPO in PNPLA3 GG variant and CC wild type models. The ratio of oxidized BODIPY (green) to reduced BODIPY (yellow) reflects the overall LPO level in each model. (A) Representative images of BODIPY 581/591 C11 staining in each PNPLA3 genotype and media condition, 40X; scale 50µm. (B) A significant increase in the ratio of OxBODIPY / ReBODIPY was observed in PNPLA3 GG variant 96-well plate LAMPS compared to the CC wild type in each media condition. Data were obtained on Day 5 with n = 4 wells from a single patient lot for each PNPLA3 genotype and plotted mean ± SEM for each media condition. Statistical significance was assessed by ANOVA with Tukey's test. p-value < 0.05 was considered statistically significant.

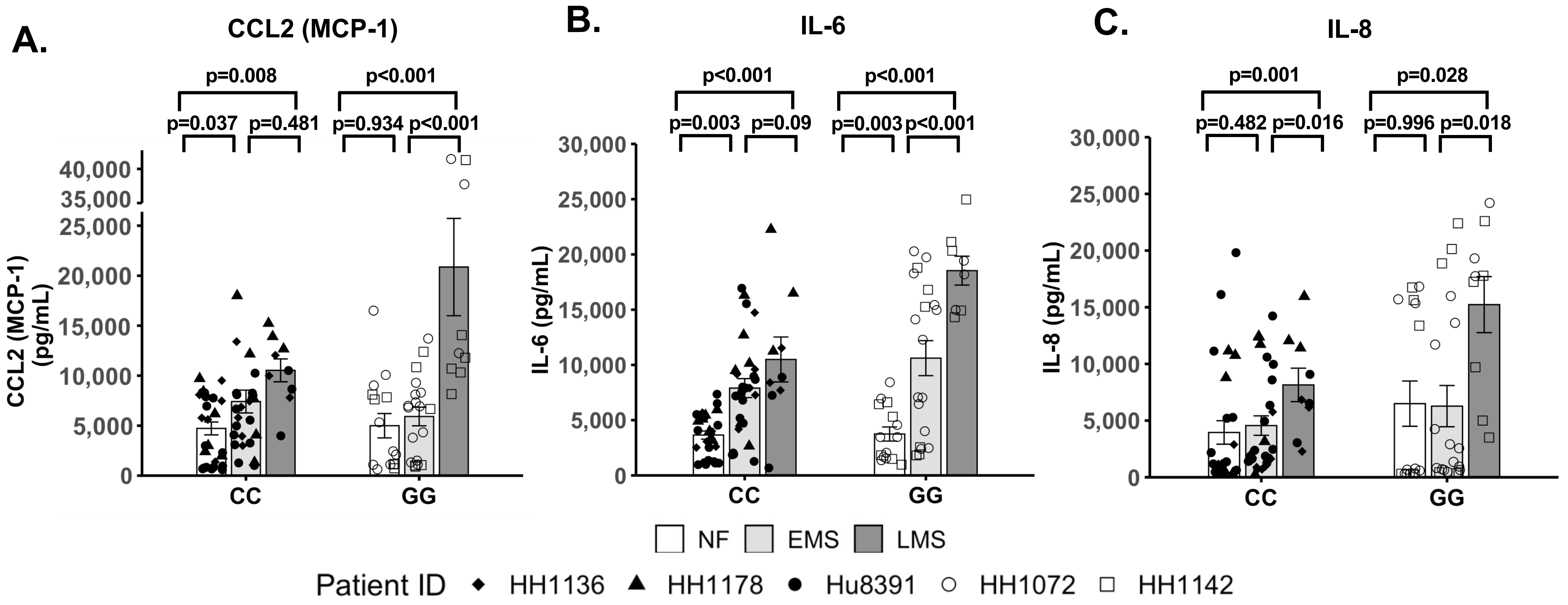


**Figure S5. Increased pro-inflammatory cytokine secretion was observed in both EMS and LMS media compared to NF medium in PNPLA3 CC and GG LAMPS demonstrating lifestyle driven inflammation.** (A-C) Significant increases in the secretion of CCL2 (A), IL-6 (B) and IL-8 (C) were observed in both EMS and LMS media compared to NF medium for each PNPLA3 genotype. While all three cytokines showed significant increases in LMS medium compared to EMS medium in PNPLA3 GG variant LAMPS (A-C), only IL-8 secretion was significantly increased in PNPLA3 CC wild type LAMPS (C) in LMS medium, consistent with recent studies demonstrating increased immune activation and inflammation associated with the PNPLA3 GG variant (7-9). Data were obtained on Day 8 with a minimum of n = 3 LAMPS from each patient lot for each condition and plotted mean ± SEM. Statistical significance was assessed by ANOVA with Tukey's test. p-value < 0.05 was considered statistically significant. These data support the analysis performed in Figure 3.

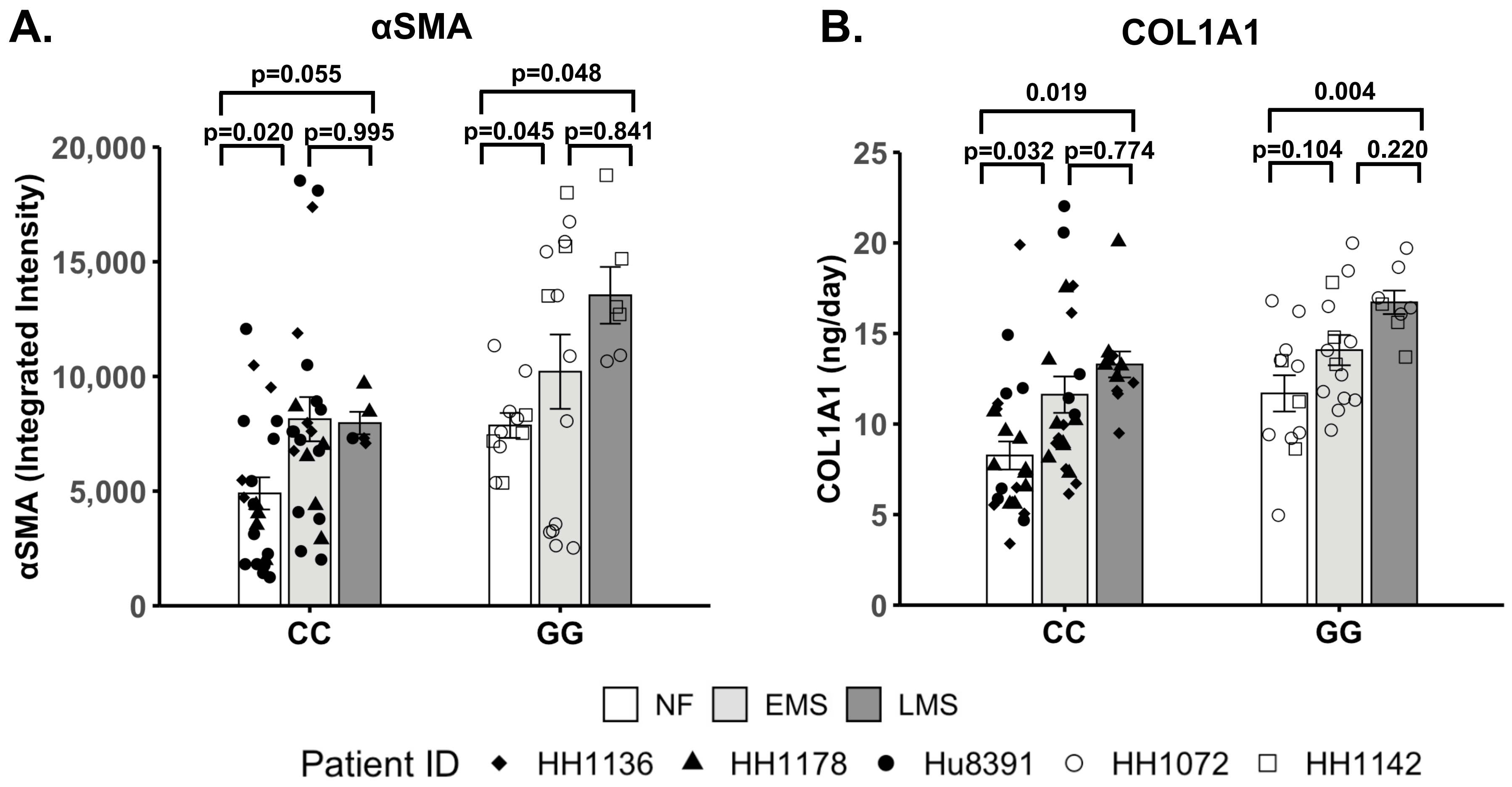


**Figure S6. Increased stellate cell activation and COL1A1 secretion are observed in both EMS and LMS media compared to NF medium in PNPLA3 CC and GG LAMPS demonstrating the impact of lifestyle in addition to the genotype.** (A) A significant increase in αSMA integrated intensity was observed between EMS and LMS compared to NF within each PNPLA3 genotype; however, no significant differences were observed between EMS and LMS media for either PNPLA3 genotype. (B) A significant increase in COL1A1 secretion was observed in LMS medium compared to NF medium for PNPLA3 genotypes. Data were obtained on Day 8 with a minimum of n = 3 LAMPS from each patient lot for each media condition and plotted mean ± SEM. Statistical significance was assessed by ANOVA with Tukey's test. p-value < 0.05 was considered statistically significant. These data support the analysis performed in Figure 4.

**Table S3. Drug binding for resmetirom used in LAMPS studies.**

| Media source | Resmetirom Concentration (μM) |
|---|---|
| Media Blank | Not detected |
| Input media (t = 0 hr) | 1.15 μM (0.500 μg/mL) |
| Chip #1 efflux (t = 72 hr) | 1.25 μM (0.545 μg/mL) |
| Chip #2 efflux (t = 72 hr) | 1.17 μM (0.511 μg/mL) |

To assess the drug binding capability of the polydimethylsiloxane (PDMS)-containing LAMPS device for resmetirom (FW = 435.22), we used perfusion flow tests and mass spectrometry analysis of flow through collected from LAMPS devices at 72 h to determine the overall effective concentration of resmetirom compared to the starting concentration (input) of drug as previously described (3,5,6).

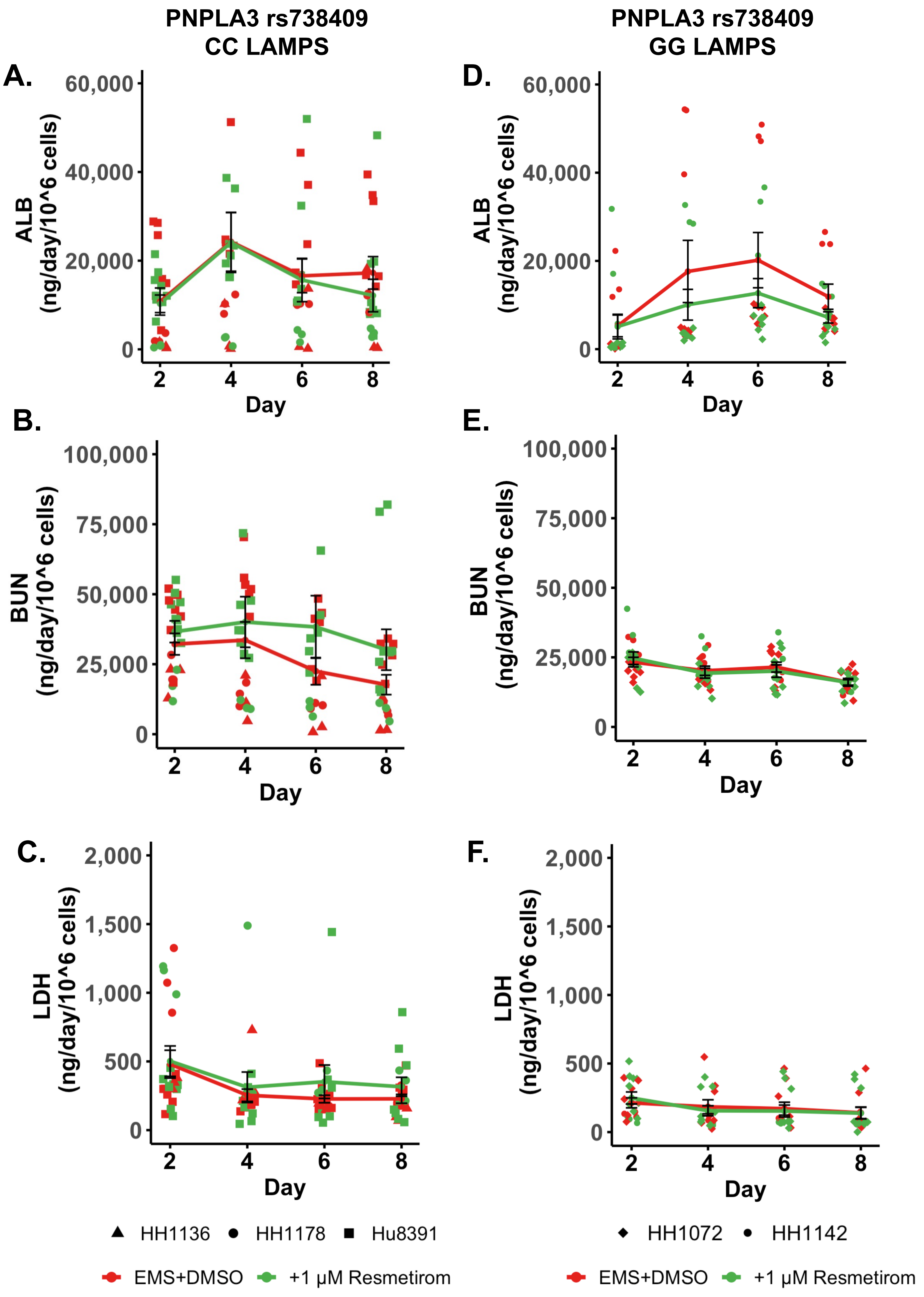


**Figure S7. Similar model functionality and cytotoxicity profiles were observed in PNPLA3-LAMPS treated with resmetirom compared to vehicle control, demonstrating that there were no significant adverse effects of resmetriom treatment.** For PNPLA3 CC wild type LAMPS (A-C) and PNPLA3 GG variant LAMPS (D-F), albumin (ALB; A and D), blood urea nitrogen (BUN; B and E) and lactate dehydrogenase (LDH; (C and F) secretion profiles were monitored to assess LAMPS functionality and cytotoxicity of 1 µM resmetirom treatment. No significant changes were observed between resmetirom treatment and vehicle control for ALB, BUN, or LDH secretion, demonstrating that drug treatment does not affect model functionality and cytotoxicity. Data were plotted mean ± SEM with a minimum of n = 3 LAMPS from each patient lot for each treatment condition. Statistical significance was assessed by ANOVA with Tukey's test for each indicated time point. p-values < 0.05 were considered statistically significant.

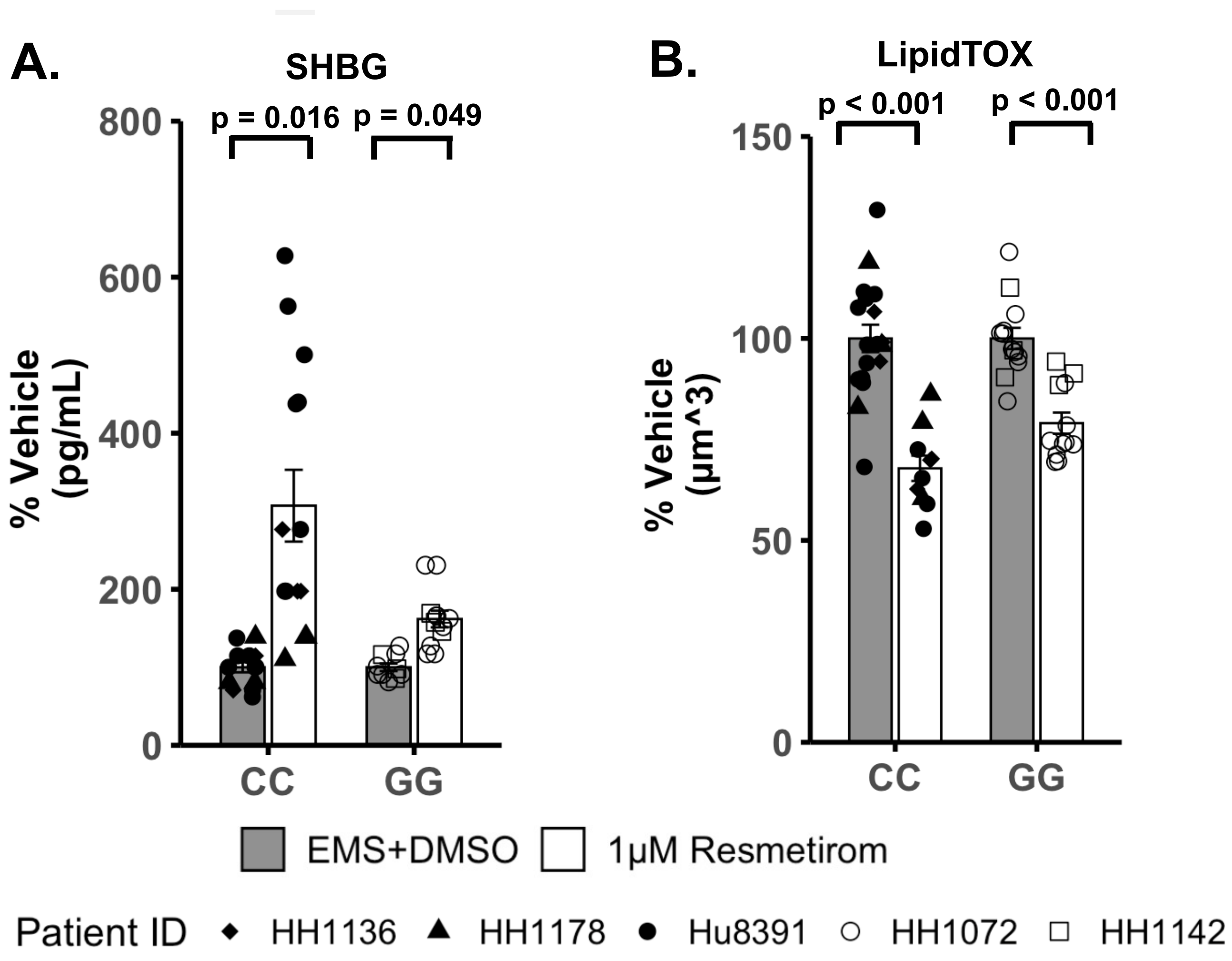


**Figure S8. Resmetirom treatment resulted in increased SHBG secretion and steatosis reduction in both PNPLA3 GG variant and CC wild type LAMPS demonstrating the pharmacodynamic effect of resmetirom treatment.** (A) Significantly increased SHBG secretion was observed in both PNPLA3 CC wild type and GG variant LAMPS with 1 µM resmetirom treatment. (B) A significant reduction in steatosis was observed in both PNPLA3 CC wild type and GG variant LAMPS with 1 µM resmetirom treatment compared to vehicle control. Data were obtained on Day 8 with a minimum of n = 3 LAMPS from each patient lot for each condition and plotted mean ± SEM. Statistical significance was assessed by ANOVA with Tukey's test. p-value < 0.05 was considered statistically significant. These data support the analysis performed in Figure 5 and are both consistent with recent clinical evidence demonstrating the pharmacodynamic effects of resmetirom (10,11).

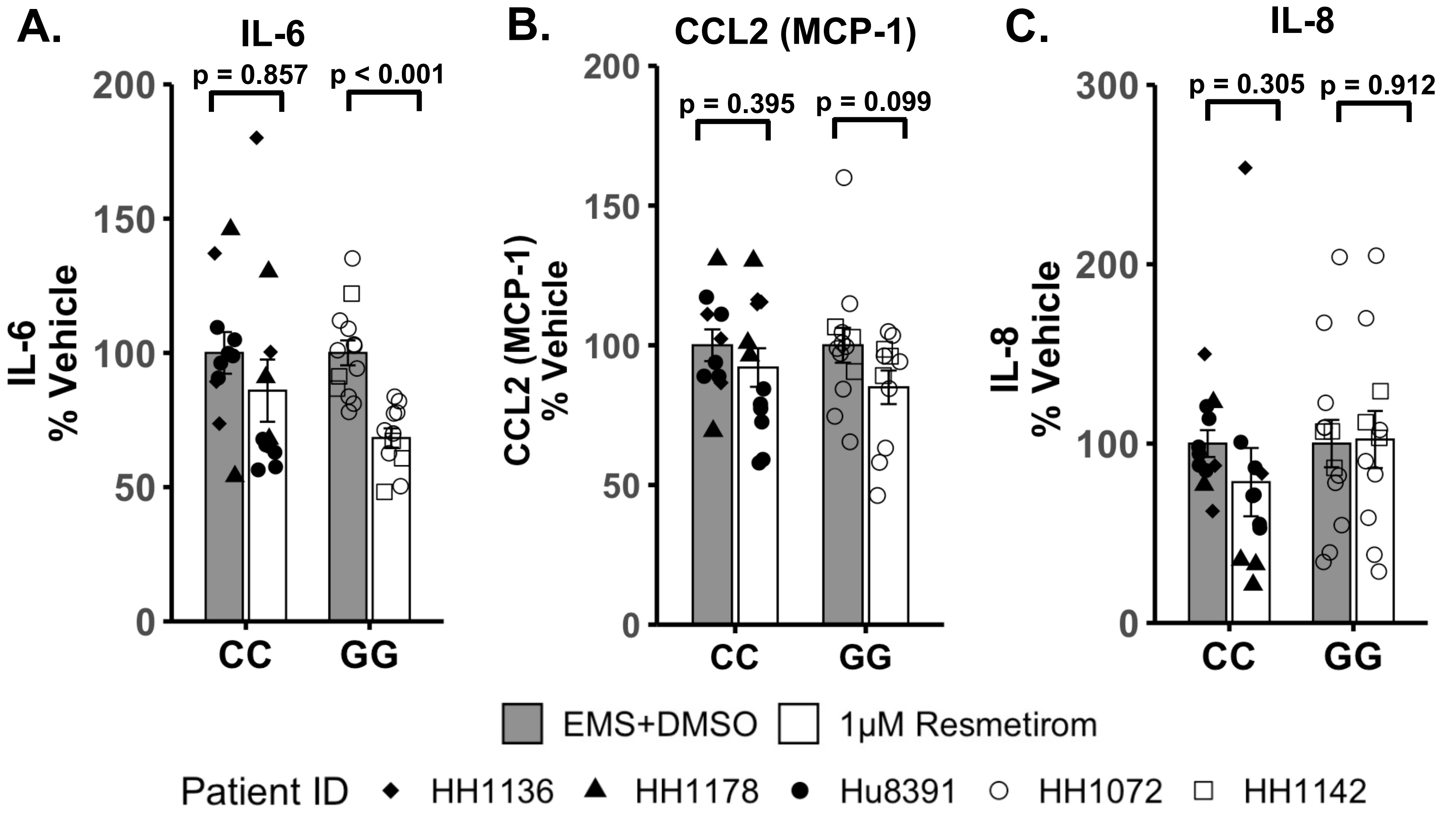


**Figure S9. Resmetirom treatment significantly reduced the secretion of the pro-inflammatory cytokine IL-6 in PNPLA3 GG variant LAMPS.** While no genotype-specific differences were observed in the reduction of secreted cytokines when PNPLA3 GG variant and CC wild type LAMPS were compared (Fig 6), Data (% of vehicle control) were obtaresmetirom treatment did result in a significant reduction of IL-6 in PNPLA3 GG variant LAMPS, but not in PNPLA3 CC wild type LAMPS when compared to their respective vehicle control (A). No significant reduction was observed for the secretion of CCL2 (B) or IL-8 (C) in either PNPLA3 LAMPS genotype when compared to their respective vehicle control.ined on Day 8 with a minimum of n = 3 LAMPS from each patient lot for each condition and plotted ± SEM. Statistical significance was assessed by ANOVA with Tukey's test. p-values <0.05 were considered statistically significant.

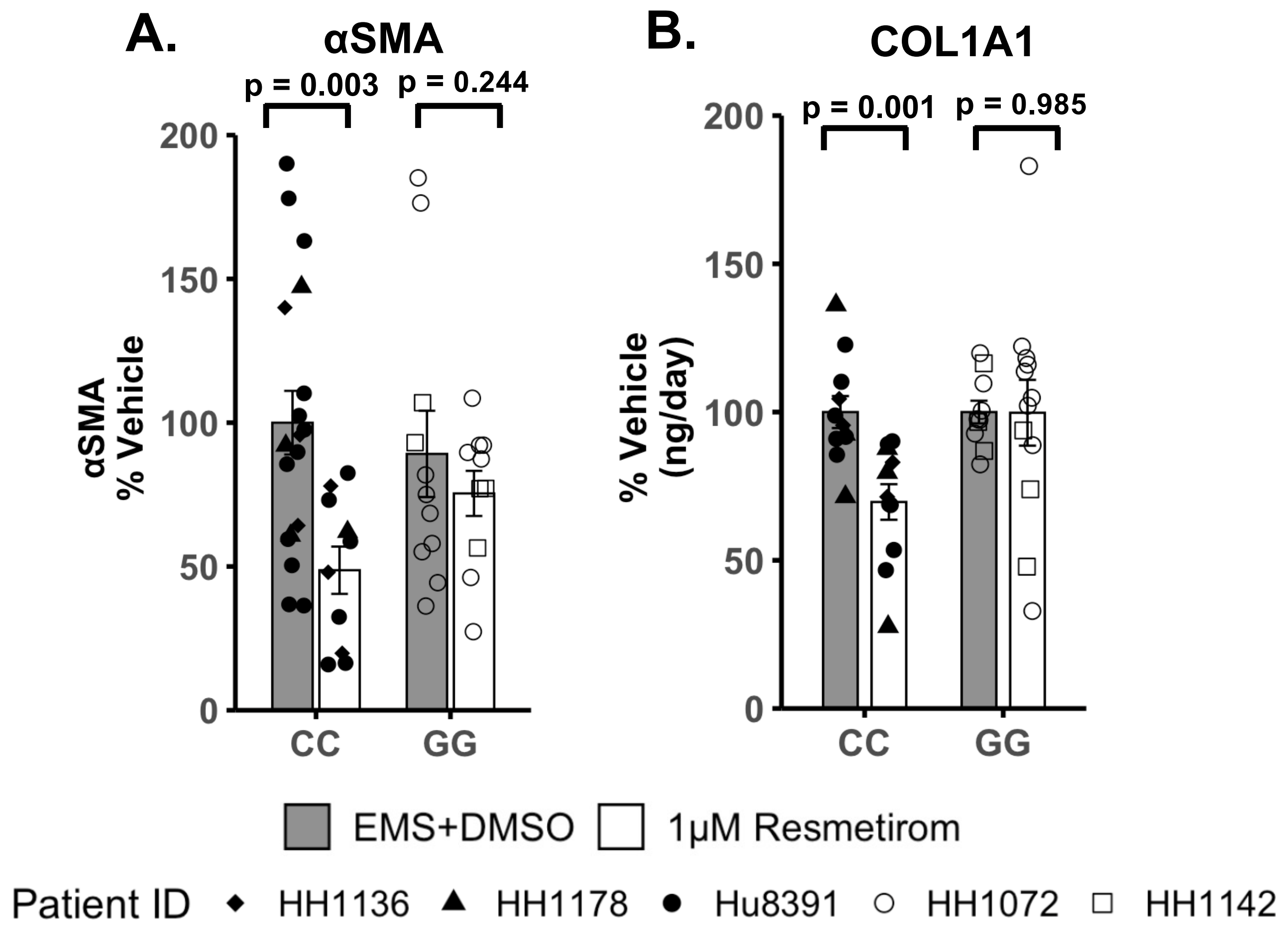


**Figure S10. Resmetirom treatment resulted in a significant reduction in stellate cell activation and COL1A1 secretion in PNPLA3 CC wild type LAMPS but not in GG variant LAMPS demonstrating a genotype-specific drug response.** (A and B) Compared to vehicle control, 1μM resmetirom treatment significantly reduced both αSMA integrated intensity (A) and the secretion of COL1A1 (B) in PNPLA3 CC wild type LAMPS, but not in GG variant LAMPS. Data were obtained on Day 8 with a minimum of n = 3 LAMPS from each patient lot for each condition and plotted mean ± SEM. Statistical significance was assessed by ANOVA with Tukey's test. p-value < 0.05 was considered statistically significant. These data support the analysis performed in Figure 7.

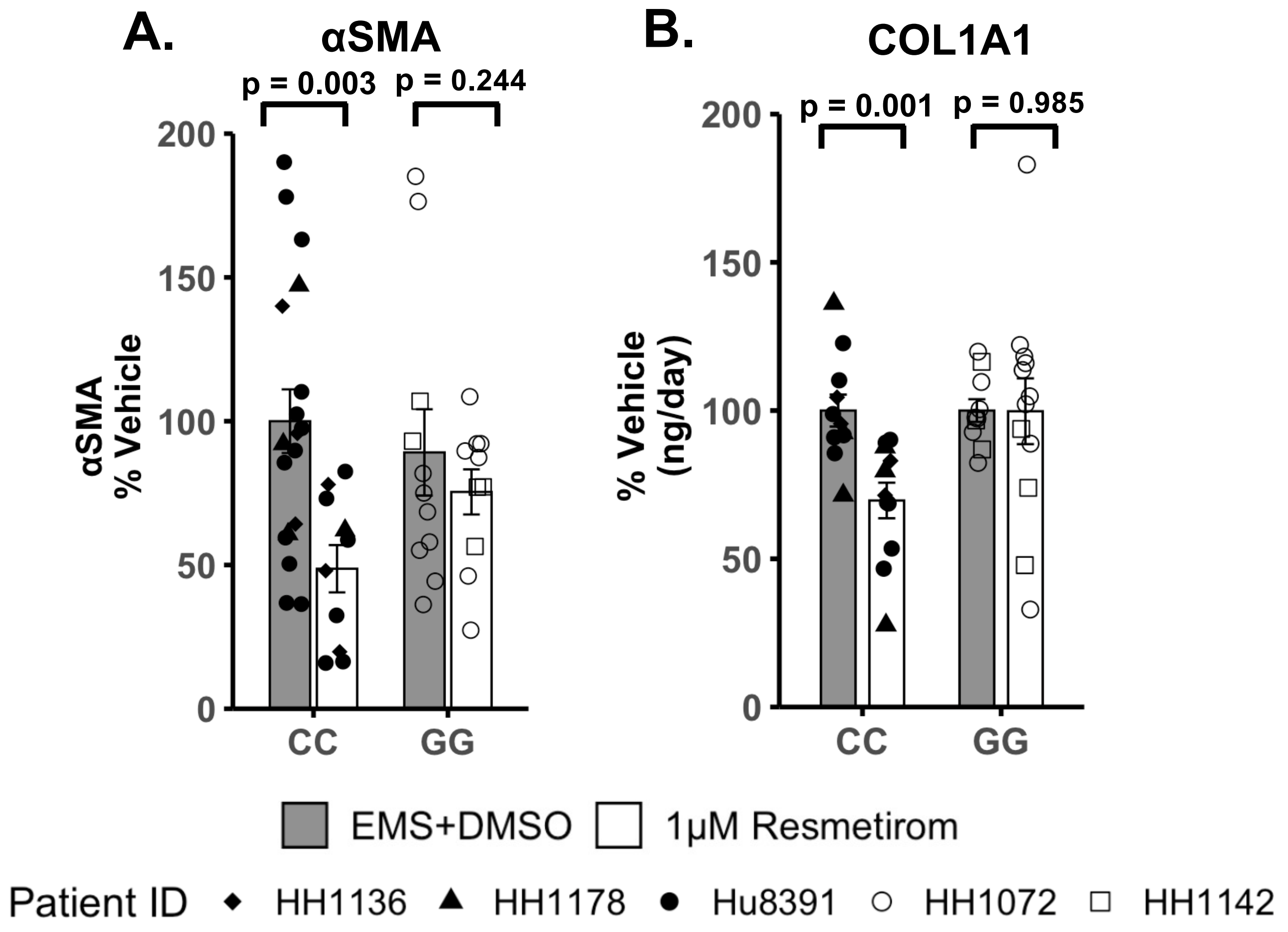


**Figure S10. Resmetirom treatment resulted in a significant reduction in stellate cell activation and COL1A1 secretion in PNPLA3 CC wild type LAMPS but not in GG variant LAMPS demonstrating a genotype-specific drug response.** (A and B) Compared to vehicle control, 1μM resmetirom treatment significantly reduced both αSMA integrated intensity (A) and the secretion of COL1A1 (B) in PNPLA3 CC wild type LAMPS, but not in GG variant LAMPS. (B) 1 μM resmetirom treatment significantly reduced COL1A1 secretion in PNPLA3 CC wild type LAMPS, while no significant difference was observed in GG variant LAMPS. Data were obtained on Day 8 with a minimum of n = 3 LAMPS from each patient lot for each condition and plotted mean ± SEM. Statistical significance was assessed by ANOVA with Tukey's test. p-value < 0.05 was considered statistically significant. These data support the analysis performed in Figure 7.
